# Structural and genetic signatures of two classes of HCV E2 neutralizing face antibodies from non-human primates immunized with a recombinant E1E2

**DOI:** 10.1101/2025.10.07.680784

**Authors:** Yen T.K. Nguyen, Fang Chen, Erick Giang, Swati Saha, Lynn A. Ueno, Christopher Chen, Corey T. Watson, Netanel Tzarum, Ian A. Wilson, Mansun Law, Robyn L. Stanfield

**Affiliations:** Department of Integrative Structural and Computational Biology, The Scripps Research Institute, La Jolla, CA 92037, USA; Department of Immunology and Microbiology, The Scripps Research Institute, La Jolla, CA 92037, USA; Department of Biochemistry and Molecular Genetics, University of Louisville School of Medicine, Louisville, KY 40202, USA; Southwest National Primate Research Centre (SNPRC) at Texas Biomedical Research Institute, San Antonio, TX 78245, USA; Skaggs Institute for Chemical Biology, The Scripps Research Institute, La Jolla, CA 92037, USA

**Keywords:** Hepatitis C virus, non-human primates, rhesus macaque neutralizing antibody, X-ray structure, antigenic region 3

## Abstract

Hepatitis C continues to be a significant public health problem despite advancements in antiviral therapeutics. To eliminate this disease, an effective vaccine against new infections and re-infections is needed. However, to date only one Hepatitis C virus (HCV) envelope protein (E1E2) immunogen, developed by Chiron Inc., has been tested in a Phase I clinical trial (ClinicalTrials.gov identifier NCT00500747). To establish a benchmark for elicitation of broadly neutralizing antibodies (bnAbs) by E1E2, we previously immunized non-human primates (NHPs) with this immunogen and isolated monoclonal nAbs that exhibit neutralization potency comparable to human nAbs. Here we show that NHP nAbs, encoded by germline genes IGHV1-138*01 and IGHV4-NL_5*01 (homologs of human IGHV1-69*10 and IGHV4-59*12, respectively), recognize a relatively E2 conserved region (neutralizing face) proximal to antigenic region 3 (AR3). These NHP AR3-targeting nAbs share highly similar binding modes to human AR3-targeting nAbs, suggesting a similarity in human and NHP immune responses to the same HCV immunogen.

## Introduction

Although anti-viral drugs to eliminate HCV infection are extremely effective ^1^, the number of HCV-infected individuals worldwide is still estimated to be over 50 million (World Health Organization, Hepatitis C: Fact sheet (2025), https://www.who.int/news-room/fact-sheets/detail/hepatitis-c). Despite treatment with direct-acting antiviral (DAA) drugs that can completely eradicate viral infection, some patients are developing liver cancer within 10 years of treatment ^2^. Thus, elimination of HCV may not always be sufficient to protect against cancer and other chronic health problems. Additionally, viral clearance via DAA does not prevent future re-infection ^3^. To prevent future infections and help eradicate HCV, an effective vaccine is needed ^4^.

A vaccine to prevent infection by the induction of bnAbs should target the viral envelope proteins E1 and/or E2 that have been shown by cryoEM to form a heterodimeric complex ^5–7^. To gain cell entry, these proteins interact with multiple receptors on the cell surface, including heparan sulfate proteoglycan, low-density lipoprotein receptor, CD81, claudin, scavenger receptor-B1, and occludin ^8–12^. The CD81 receptor binding site on E2 serves as a key target for numerous nAbs ^13–18^ and is part of the E2 neutralizing face that features highly conserved and hydrophobic sites, including antigenic site 412 (AS412, aa 412-420), AS434 (aa 434-446), and the discontinuous epitope antigenic region 3 (AR3) ^17,18^. The AR3 comprises multiple structural domains, including the E2 front layer (FL, aa 421-459), CD81 binding loop (CD81BL, aa 518-535), and back layer (BL, aa 614-622) ^17,18^. Of particular interest, AR3-targeting Abs exhibit broad neutralization activity against multiple HCV genotypes, suggesting that a broadly protective vaccine is possible ^13,14,19,20^. Furthermore, crystal structures of E2 cores in complex with Fabs ^21^ or the CD81 long extracellular loop ^22,23^ have revealed that structural flexibility exists within AR3 regions, such as the FL and CD81BL. Whether this flexibility will aid, or hinder design of potential immunogens is not clear.

A large number of human bnAbs have been isolated from HCV-infected participants. Thus far, the majority of structurally characterized bnAbs utilize heavy chains (HCs) derived from the IGHV1-69 germline family and block binding to CD81 by recognition of E2 AR3 ^16,20,24–26^; however, neutralizing antibodies (nAbs) that recognize other epitopes and are encoded by other germline families are also being isolated and studied ^27^. The human IGHV1-69 germline gene is unique in that its codes for hydrophobic residues at the tip of CDRH2 that have been often observed to bind to hydrophobic surfaces on the HCV E2 antigen. Many IGHV1-69 encoded antibodies also target human viruses including influenza and HIV ^25,28,29^.

Until controlled human infection models are approved and available ^30,31^, the rhesus macaque (RM) is of interest as a model for antibody elicitation. While macaques cannot be infected with HCV, their immune system is highly similar to that of humans ^32–37^, and macaque Abs elicited by immunization can be compared to their human counterparts. However, despite the growing availability of NHP genomic databases ^38,39^, the RM IG loci remain poorly understood because of their genomic complexity and incomplete assembly ^40–42^. The identification of individual germline genes for each tested NHP animal can ensure the accurate assignments of identified Abs to specific germline genes/alleles and estimation of SHM ^38,42,43^. This process is critical for tracing Ab affinity maturation pathways in vaccinated NHP as a means to identify parallels in human-NHP genetics ^36,37,44,45^.

Surprisingly, the only human vaccine trial carried out to date using a hepatitis envelope protein as immunogen was initiated in 2003 by Chiron (ClinicalTrials.gov identifier NCT00500747) ^46^, where a recombinant, full-length, genotype 1a E1E2 envelope protein was used as immunogen with MF59 as adjuvant. This vaccine advanced to a Phase I trial, but cross-neutralizing responses were produced in only a few vaccine recipients ^47^. We previously obtained and analyzed serum samples from that trial ^44^; however, no samples were available from which Abs could be isolated. Thus, to better understand this study, we previously immunized four RMs with the Chiron E1E2 immunogen (Fig. 1a) and found that both animals and humans had similar Ab responses ^44,45^. Although the immune sera are largely strain-specific, monoclonal nAbs can be isolated from RM plasmablasts and memory B cells ^44,45^. Many of these nAbs target AR3 and are mostly derived from germline IGHV1-138*01_S6073 (KIMDB nomenclature ^38^) that is homologous to the human IGHV1-69*10 gene (94.6% DNA sequence identity) ^44,45^ (Fig. S1).

**Fig. 1:**
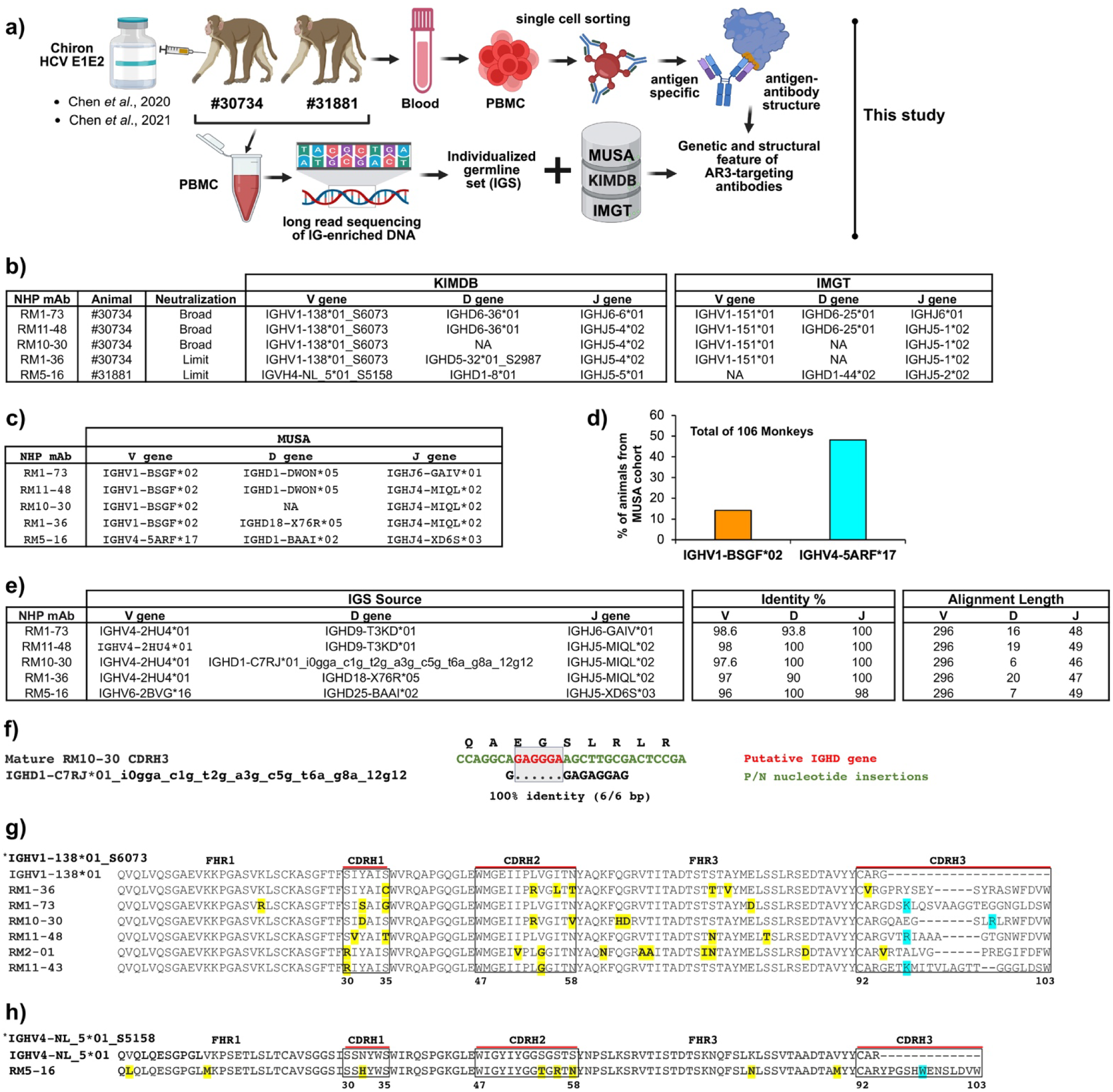
Overall genetic analysis of AR3-targeting nAbs from NHP. **a** Overview: sample collection, single-cell sorting, and individualized germline set (IGS) generation. Two rhesus monkeys (RMs #30734 and #31881) were vaccinated with Chiron HCV E1E2, as previously described ^44,45^. Their peripheral blood mononuclear cells (PBMCs) were collected, followed by single-cell sorting to isolate antigen-specific B cells and mAbs. In parallel, targeted single-molecule real-time (SMRT; Pacific Biosciences) long-read sequencing was performed on PBMC-derived genomic DNA from these same vaccinated RMs, generating an IGS. The Figure was created with BioRender.com. **b, c** General information of five RM nAbs that were isolated from RMs #30734 and #31881. Their genetic germline allele feature is assigned by KIMDB ^38^, IMGT ^39^, and MUSA database ^48^. NA is not identical. The germline gene assignments were determined by three criteria: (1) matching of V and J gene usage, (2) identical CDR3 length, and (3) CDR3 nucleotide sequence homology >80%. **d** Frequency of IGHV1-BSGF*02 (orange) and IGHV4-5ARF*17 (cyan) germline alleles. The Y-axis represents the detection frequency of these germline alleles among 106 monkeys from the MUSA database ^48^. **e**. The V, D, and J germline allele assignment by our IG long-read sequencing and IGS. **f** IGS reveals the closest putative IGHD gene for RM10-30 Ab. Sequence alignment was performed between mature Ab and identified IGHD germline gene by IGS. Sequence of IGHD gene and P/N nucleotide insertion region are in red and green, respectively. Dots denote the matching nucleotides with mature Ab sequence. **g, h** HC sequence alignment of RM Abs with IGHV1-138*01 and IGHV4-NL_5*01 germline alleles. The variant residues of RM Abs in comparison with encoded germline alleles are in bold and shaded in yellow. The CDRH3 basic, long side chain residues of IGHV1-138*01 encoded RM Abs are shaded in red. The key aromatic residue of CDRH3 RM5-16 is shaded in cyan. CDR1-3 length and sequence are defined according to Kabat numbering.

To gain deeper insight into the conserved properties of AR3-targeting nAbs, we performed genetic, structural, and functional characterization of five AR3-targeting nAbs from RMs immunized with the Chiron E1E2 vaccine ^44,45^. We also confirmed their verified germline sequences directly through targeted long-read IG loci genomic sequencing of the two study animals (#30734 and #31881) to verify the genetic features of these RM AR3-targeting class Abs. Four of these nAbs (RM1-73, RM11-48, RM10-30, and RM1-36) utilize the same IGHV gene segment, IGHV1-138*01_S6073, with limited SHM and exhibit a very similar E2 binding mode as Abs isolated from human elite neutralizers ^20^. NAb RM5-16 is derived from a different germline gene, IGHV4-NL_5*01_S5158, and recognizes a similar epitope footprint but with a different binding approach angle to E2. These studies show that the NHP immune system can mimic that of humans in response to HCV immunogens but can also recognize the conserved AR3 epitope (within neutralizing face) in different ways using different germline genes.

## Results

### Genetic features of vaccine-induced AR3-targeting Abs from RM

Previously, we reported the isolation of 100 HCV-specific mAbs from two Chiron E1E2-immunized RMs: 56 from RM#3074 and 44 from RM#31881 (Fig. 1a) ^44,45^. Of these, 24 mAbs exhibited neutralizing activity (15 from RM#3074 and 9 from RM#31884) ^45^. Twelve of these nAbs (including RM1-73, RM11-48, and RM10-30) demonstrated potent cross-neutralizing activity (classified as bnAbs), 10 (including RM1-36 and RM5-16) showed limited breadth, and the remaining two were strain-specific ^45^.

Germline gene usage of five RM AR3-targeting nAbs (RM1-73, RM11-48, RM10-30, RM1-36, and RM5-16) studied was initially assigned using KIMDB ^38^ and IMGT ^39^ (Fig. 1b). More recently, we searched a data set derived from sequencing of 106 macaques (Macaque United Set of Alleles, MUSA) ^48^, which was compiled from combined genomic and repertoire sequencing. These data are searchable at: https://vdjbase.org/ reference_book/Rhesus_Macaque. Using the MUSA data and naming system, RM AR3-targeting nAb germline genes were assigned as IGHV1-BSGF*02 and IGHV4-5ARF*17 (Fig.1c). The IGHV1-BSGF*02 and IGHV4-5ARF*17 germline alleles were detected in 14% (15/106) and 48% (51/106) of macaques in the MUSA group, respectively ((Fig. 1d). None of the databases searched contained an IGHD germline allele aligning with RM10-30 while no corresponding IGHD germline gene was found for RM1-36 or IGHV germline gene for RM5-16 in the IMGT database (Fig. 1b, c).

To confirm the authentic germline alleles of these five RM nAbs, we next employed targeted single molecule real-time (SMRT; Pacific Biosciences) long-read sequencing from PBMC-derived genomic DNA of the same vaccinated RMs (#30734 and #31881) (Fig. 1a; see Materials and Methods for details). These data allowed us to generate haplotype-resolved assemblies spanning the IG loci for both animals (#30734 and #31881), from which we conducted comprehensive curation of their germline IG alleles using Digger ^49^. Next, we performed multiple sequence alignment (MSA) of the identified Ab sequences and their corresponding best-matched germline alleles from previous databases, as well as the curated germline annotation of each animal. As a result, the direct targeted IG long-read sequencing and individualized germline set (IGS) confirmed the expected germlines allele for these five RM AR3-targeting nAbs. Specifically, RM1-73, RM10-30, RM11-48, and RM1-36 arise from IGHV4-2HU4*01 (identical to IGHV1-138*01_S6073), while RM5-16 is derived from IGHV6-2BVG*16 (identical to IGHV4-NL_5*01_S5158) (Fig. 1e and Fig. S1a). RM #30734 appears homozygous to IGHV4-2HU4*01, whereas RM #31881 is also homozygous to IGHV6-2BVG*16.

IGS identified the IGHD germline alleles for RM1-73, RM11-48, RM1-36, and RM5-16, each showing 100% DNA identity to counterparts in the other databases (Fig. 1e and Fig. S1b). Notably, the IG long-read sequencing data allowed us to find the closest putative IGHD gene for RM10-30 as IGHD1-C7RJ*01_i0gga_c1g_t2g_a3g_c5g_t6a_g8a_12g12, which showed a 6 nucleotide base pair (bp) match to the mature Ab sequence (Fig. 1f). Comparison with the putative IGHD germline allele thus shows conservation as well as variation (Fig. 1f); thus, whether it best represents the underlying germline gene for this Ab may require additional data analyses. In addition, IGS confirmed that the IGHJ germline alleles for these five RM AR3-targeting nAbs are also completely identical at the nucleotide level to corresponding germline alleles found in other databases (Fig. S1c).

Our RM germline sequences revealed consistently low SHM rates for the IGHV of these five RM AR3-targeting nAbs of 1.4-4% at the nucleotide level (Fig. 1e). The RM1-73 Ab has the lowest SHM (only 1.4%) and the longest CDRH3 of 25 aa (Fig. 1e, 1g). RM1-73, RM11-48, RM10-30, and RM1-36, which were all isolated from the same animal, use the same IGHV (Fig. 1e, 1g, and Fig. S1a). RM1-73 and RM11-48 also used the same IGHD gene, while RM11-48, RM10-30, and RM1-36 used the same IGHJ (Fig. 1e and Fig. S1b). In contrast, these RM Abs used different IGHD and IGHJ genes from those of previously described RM2-01 and RM11-43 Abs ^44,45^. RM5-16, isolated from a different animal, used distinct IGHV, D, J germline genes (Fig. 1e, 1h, and Fig. S1a).

To compare rhesus and human homologs, we also performed MSAs of these IGHV alleles at both nucleotide and amino acid levels, incorporating their five closest human alleles to illustrate evolutionary similarity (Figs. S1d, S1e). From these alignments, we identified IGHV1-69*10 (94.6 % DNA identity and 90.8 % protein identity, L at codon 54) as the closest human counterpart to IGHV1-138*01_S6073, although differences to various IGHV1-69 alleles were small (0.7% at the DNA and 0 % at protein level). IGHV4-NL_5*01_S5158 aligned most closely with IGHV4-59*12 (93.6% DNA and 92.9 % protein identity), but again with small differences to other V_H_4-59 alleles (Fig. S1d, S1e). Henceforth, we will use the KIMDB germline identifiers for clarity and abbreviate IGHV1-138*01_S6073 to IGHV1-138*01, and IGHV4-NL_5*01_S5158 to IGHV4-NL_5*01.

### Overall structure of RM IGHV1-138*01 class Abs and complexes with HK6a E2c3

We determined structures for the four RM IGHV1-138*01 class Fabs RM11-48, RM10-30, RM1-73, and RM1-36 in complex with the E2 core domain (E2c3) from HCV isolate HK6a, as well as an unliganded structure for Fab RM1-73 (Fig. 2, Fig. S2, and Tables S1, S2). The complex structures, ranging from 2.6-2.9 Å resolution, revealed that all four RM Fabs recognize the canonical AR3 region of HK6a E2c3 (Fig. 2b), consistent with previous epitope mapping data ^45^. The Fab binding to E2c3 arises mainly from the HC, which accounts for 85% (RM1-73), 65% (RM11-48), 79% (RM10-30), and 90% (RM1-36) of the total buried surface area (BSA) on E2c3 (Fig. 2c left panel). This HC domination is also observed in human anti-HCV IGHV1-69 class Abs and two previously reported RM IGHV1-138*01 class Abs RM2-01 and RM11-43 (Fig. 2c left panel) ^45^. Most of the interactions are from CDRH3, which contributes 45-59% of the total BSA of the HC (Fig. 2c right panel).

**Fig. 2:**
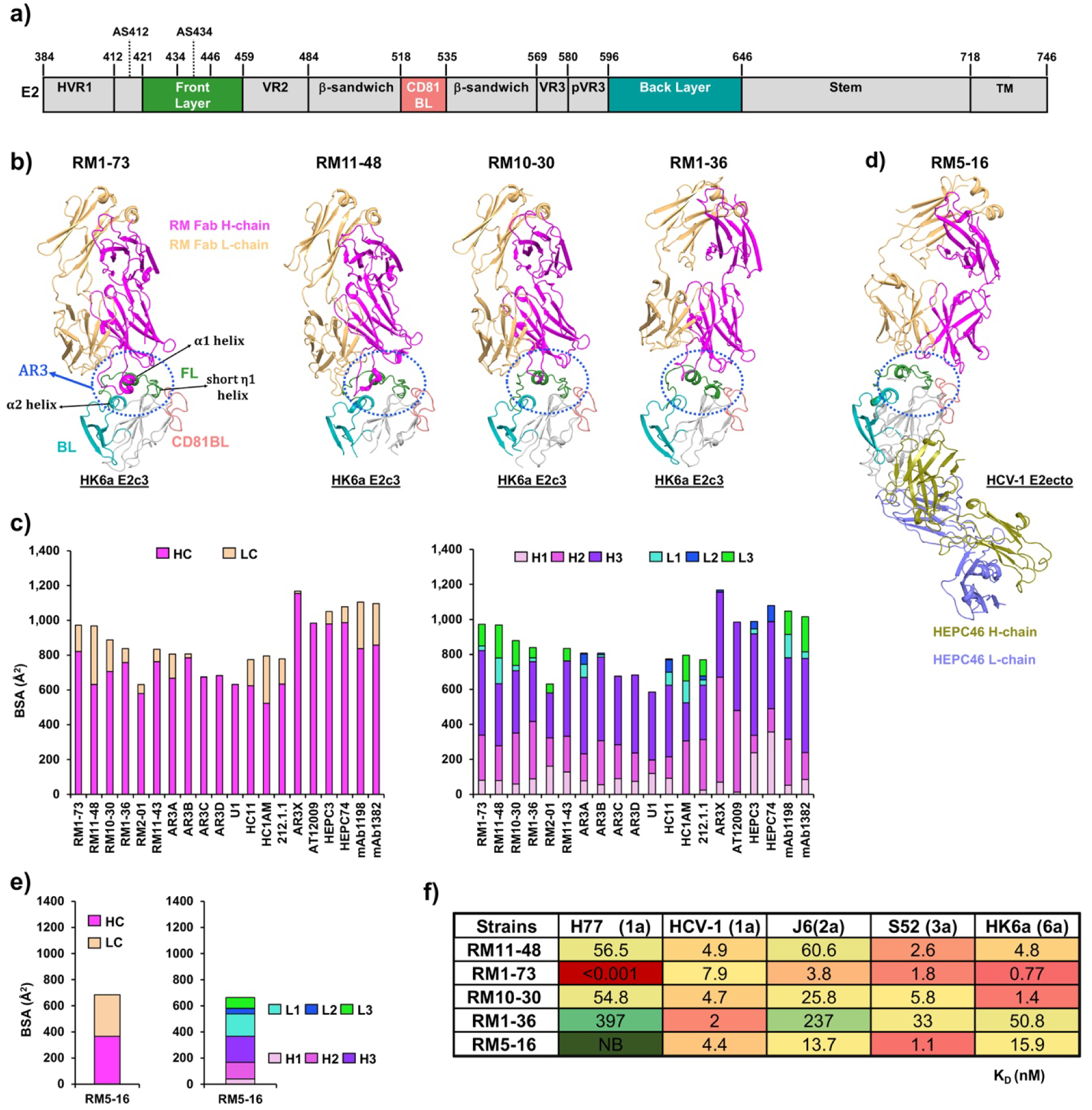
Overall structure of E2 with RM IGHV1-138*01 (RM11-73, RM11-48, RM10-30, RM1-36) and IGHV4-NL_5*01 RM5-16 Abs. **a** E2 subregions are colored as in schematic, including HVR1 (aa 384-420), AS412 (aa 412-420), front layer (FL, aa 421–458), AS434 (aa 434-446), variable region 2 (VR2, aa 459–483), β-sandwich core (aa 484–517 and 535–568), CD81 binding loop (CD81BL, aa 518–534), variable region 3 (VR3, aa 569–579), post-variable 3 region (pVR3, aa 580–595), back layer (BL, aa 596–645), stem (aa 646–717), transmembrane (TM, aa 718-746). E2 FL, CD81BL, and BL subregions are colored in green, salmon, and teal, respectively with the remainder of HK6a E2c3 colored grey. **b** The crystal structures reveal all IGHV1-138*01 RM Abs target AR3 region (AR3: FL, N-terminus (aa 421-436), helix α1 C-terminus (aa 438-443), middle of β-sandwich (aa 504-508), CD81BL (aa 528-531), and helix α2 of BL (aa 614-622)). AR3 is highlighted in a blue dotted circle. HC and LC of RM Abs are shown in magenta and light orange, respectively. **c** Buried surface area (BSA) analysis of human IGHV1-69, and RM IGHV1-138*01 HC and LC CDRs indicate the HC makes most of the E2 interactions. Structures of E2 with RM1-73 (PDB ID 9MNT), RM11-48 (9MNU), RM10-30 (9MNQ), RM1-36 (9MNS), AR3A (6BKB), AR3B (6BKC), AR3C (6UYD), AR3D (6BKD), AR3X (6URH), U1 (6WO3), HC11 (6WO4), HC1AM (6WOQ), 212.1.1 (6WO5), AT12009 (7T6X), mAb1198 (7RFB), mAb1382 (7RFC), HEPC3 (6MEI), HEPC74 (6MEH), RM2-01 (7JTF), and RM11-43 (7JTG) are used for BSA analysis. Left panel, BSA of HC and LC are in magenta and wheat, respectively. Right panel, BSA of CDRH1 (H1), CDRH2 (H2), CDRH3 (H3), CDRL1 (L1), CDRL2 (L2), and CDRL3 (L3) are in pink, magenta, purple, cyan, blue, and green, respectively. **D** Crystal structure of IGHV4-NL_5*01 RM5-16 with HCV-1 E2ecto and HEPC46. HC and LC of HEPC46 are shown in deep olive and slate, respectively. **e** BSA analysis of RM5-16 HC/LC CDRs upon binding to E2ecto. **f** BLI binding of all RM Fabs with HCV E2 proteins from different isolates and genotypes. The strongest binding is in red and no binding (NB) in dark green. The gradient yellow-green color indicates the mid-range binding affinity.

Epitope analysis revealed all four RM Fabs recognize largely overlapping epitopes at the central hydrophobic groove within the AR3 region, contacting a cluster of hydrophobic residues including the E2 FL N-terminus (I422, T425, and L427), α1 helix (I438, T439, L441, and F442), C-terminus (Y443 and A444), β-sandwich (P505), CD81BL (W529), and BL (P612 and Y613) (Fig. 3 left panel). Among these residues, T425, L427, L441, Y443, P505, W529, P612, and Y613 are highly conserved across 6 HCV genotypes (Fig. S3), suggesting all RM Abs recognize a conserved hydrophobic core of E2. RM11-48 engages additional hydrophobic residues (V447, V622, and L626) adjacent to the core hydrophobic groove, whereas RM1-73 uniquely contacts the AS412 region (N417-W420) (Fig. 3 left panel).

**Fig. 3:**
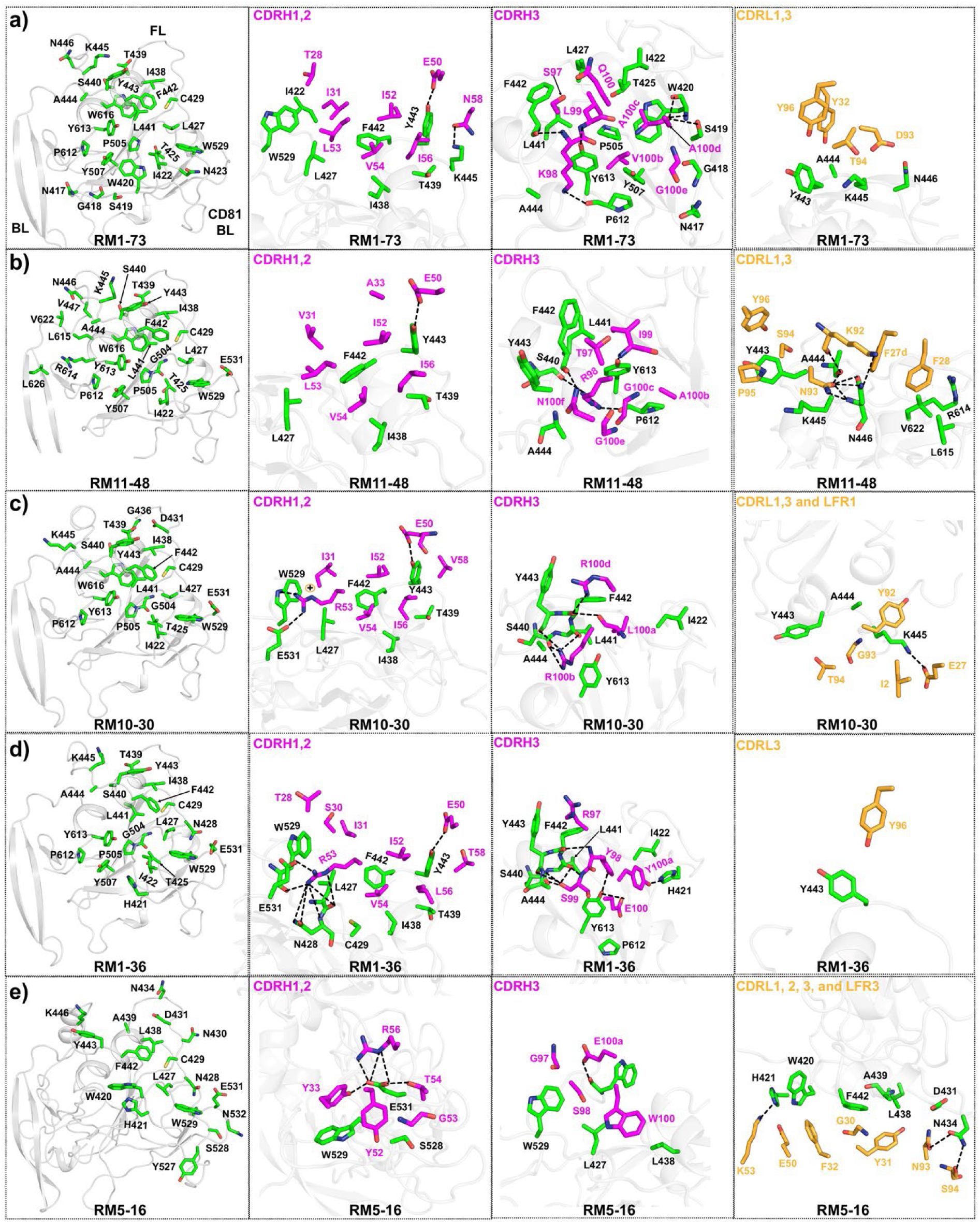
Interaction of RM Abs with HCV E2 proteins. **a**-**e** Left panel, E2 epitope residues (with BSA > 0 Å^2^ according to PISA analysis) are in green sticks, and E2 protein backbones are shown in grey tubes. Mid to right panel: E2 (green) detailed interactions with the RM Ab HC (magenta) and LC (gold). The E2 and RM Ab residues with BSA > 0 Å^2^ and within 3.9 Å are labeled and shown as sticks. The H-bonds and charged interactions are in dashed black lines. The π-cation interaction is highlighted in an orange circle.

**RM1-73**: Fab RM1-73 undergoes a large conformational change upon binding E2, where CDRH3 changes from a loop conformation in the unliganded Fab to a helical conformation when bound to E2 (Fig. S2a), with a Cα root-mean-square deviation (RMSD) of 5.3 Å for CDRH3 residues C92-W103 in bound and unbound Fab structures. In the E2-RM1-73 complex structure, the RM1-73 HC and LC both interact with E2 (Figs. 2c, 3a), burying 972 Å^2^ of surface on the E2 (821 Å^2^ for HC and 151 Å^2^ for LC). CDRH1 forms many hydrophobic interactions with E2 core hydrophobic groove residues I422, L427, F442, and W529 (Fig. 3a). Whereas CDRH2 of previously characterized Abs RM2-01 interacts only with the C-terminus of the FL α1 helix (aa 439-443) within the core hydrophobic groove ^45^, CDRH2 of RM1-73 retains this footprint and makes additional contact towards the FL N-terminus L427 and C-terminus K445 (Fig. 3a). Two hydrogen bonds (H-bonds) are formed between CDRH2 E50 and N58 with E2 Y443 and K445, respectively (Fig. 3a and Table S3). Compared to CDRH1-2, the CDRH3 loop binds lower in the core hydrophobic groove and extends toward AS412 (G418, S419, and W420), FL C-terminal tail A444, β-sandwich P505, and the BL (P612 and Y613) (Fig. 3a and Table S4). CDRH3 K98 H-bonds with E2 F442 and P612, while CDRH3 L99 H-bonds with Y613 (Fig. 3a and Table S3). CDRH3 A100c backbone H-bonds with conserved AS412 S419 and W420 (Fig. 3a, S3, and Table S3). The only contacts from the LC arise from CDRs L1 and L3 (Fig. 3a and Tables. S3, S4), where CDRL1 Y32 contacts E2 FL A444 and CDRL3 D93/T94 interact with the FL helix α1 Y443, K445, and N446.

**RM11-48**: RM11-48 binds E2 with a BSA of 968 Å^2^ (632 Å^2^ for HC and 336 Å^2^ for LC) (Fig. 2c). The CDRH1-2 loops of RM11-48 bind E2, recognizing a similar epitope as those of RM1-73 (Fig. 3b). Like RM1-73, RM11-48 CDRH3 binds into the AR3 hydrophobic groove (Fig. 3b and Table S4). CDRH2 E50 H-bonds with E2 Y443 and CDRH3 R98 engages in a H-bond network with E2 FL S440 and BL P612; I99 H-bonds to Y613, and N100f H-bonds to F442 (Fig. 3b and Table S3). Among the four RM Fabs, the LC of RM11-48 forms the most extensive interactions with E2 (Fig. 3b and Tables S3, S4). CDRL1 F27d and F28, as well as CDRL3 N93, P95, and Y96 create a hydrophobic cluster that interacts with another hydrophobic patch formed by the E2 FL C-terminus (Y443 and A444) and BL C-terminus (L615 and V622) (Fig. 3b and Table S4). CDRL3 K92 and N93 form H-bonds with E2 A444 and N446 (Fig. 3b and Table S3).

**RM10-30**: RM10-30 recognizes E2 with 889 Å^2^ of BSA (706 Å^2^ for HC and 182 Å^2^ for LC) (Fig. 2c). RM10-30 CDRH1-2 bind into a slightly wider E2 AR3 hydrophobic patch as compared to that bound by RM11-48 (Fig. 3c and Table S4). CDRH1 T28 is within H-bonding distance of the N423 glycan (Table S3). CDRH2 E50 also H-bonds with E2 helix α1 Y443, while R53 forms a salt bridge with CD81BL E531 and a π-cation interaction with W529 (Fig. 3c and Table S3). CDRH3 L100a binds in the lower hydrophobic pocket bordered by E2 I422, L441, F442, and Y613 (Fig. 3c and Table S4) while CDRH3 R100b and R100d form several H-bonds with E2 FL helix α1 S440, L441, F442, and Y443 (Fig. 3c and Tables S3, S4). CDRH3 R100b forms π-π interaction with E2 Y443 (Fig. 3c and Table S4). CDRL3 is involved in hydrophobic interactions with E2 FL helix α1 residues Y443 and A444 while CDRL1 E27 forms a charged interaction with E2 C-terminal residue K445 (Fig. 3c and Tables S3, S4). The LC framework LFR1 I2 makes a van der Waals (vdW) interaction with E2 K445 (Fig. 3c and Table S4).

**RM1-36**: RM1-36 buries 839 Å^2^ of surface area (758 Å^2^ for HC and 81 Å^2^ for LC) in its complex with E2 (Fig. 2c). CDRH1 makes hydrophobic interactions with E2 FL helix F442 and CD81BL W529 (Fig. 3d and Table S4). CDRH2 binds to the same hydrophobic patch as CDRH2 from RM10-30 (Fig. 3d and Table S4). As for the other three RM IGHV1-138*01 Fabs, CDRH2 E50 forms a H-bond with E2 α1 helix Y443 (Fig. 3d and Table S3). CDRH2 R53 engages in polar and charged interactions with E2 FL N428 and CD81BL E531, H-bonds with FL L427/N428 and CD81BL W529 (Fig. 3d and Table S3, S4). CDRH3 also occupies the AR3 core hydrophobic groove (Fig. 3d and Table S4). CDRH3 S99 forms several H-bonds with E2 FL S440, L441, Y443, and A444, while Y98 H-bonds with FL F442 and BL Y613, and E100 also H-bonds to BL Y613 (Fig. 3d and Table S3). Additionally, CDRH3 Y100a H-bonds with E2 FL N-terminus H421 (Fig. 3d and Table S3). The RM1-36 LC has little contact with E2, with only CDRL3 Y96 forming a hydrophobic interaction with FL Y443 (Fig. 3d).

### Characterization of RM IGHV4-NL_5*01 class Ab RM5-16 and structure with HCV-1 E2ecto

To investigate the structural basis of RM IGHV4-NL_5*01 class Ab recognition, crystal structures of RM5-16 Fab, unliganded and with the HCV-1 strain E2 ectodomain (E2ecto, aa 384-645), were determined at 1.46 and 3.39 Å resolutions, respectively (Figs. 2d, S2b, and Tables S1, S2). RM5-16 also recognizes AR3 of E2ecto (Fig. 2d), consistent with previous epitope mapping ^45^. The overall RM5-16 structures for unliganded and ligand-bound RM5-16 are highly similar, with a Cα RMSD of 0.2 Å calculated for variable domain (residues variable VH 1-113 and VL 1-110). Unlike RM1-73, there is essentially no difference in the CDR regions of regions of bound versus unbound RM5-16, with RMSD of only 0.1 Å for CDRH3 (residues C92-W103) (Fig. S2b). Unlike the four RM IGHV1-138*01 class Abs, the HC and LC of RM5-16 contribute more equally to E2 binding, accounting for 54% (368 Å^2^) and 46% (317 Å^2^) of the total BSA on E2ecto, respectively (Fig. 2e). CDRH3 of RM5-16 remains the largest contributor to E2ecto binding (57% of total HC BSA) (Fig. 2e). The epitope of RM5-16 overlaps to some extent with those of RM IGHV1-138*01 class Abs but also includes contacts to the top of the E2ecto FL loop (aa 431-439) and CD81BL (aa 528-529) (Fig. 3e). Although RM5-16 has an N-linked glycosylation sequon at HC N81 and fully exposed, we observe no electron density for a glycan at that position in both unliganded and ligand-bound RM5-16 structures.

The resolution of this complex structure is relatively low (3.39 Å), limiting precise H-bond assignment; however, the high-resolution Fab structure (1.46 Å) and previously determined higher resolution E2ecto structures aided as templates during our model building. H-bonds described here are within H-bonding distance in our model. Within the HC, the RM5-16 CDRH1-2 loops contact only E2 CD81BL (Fig. 3e). CDRH1 Y33 is within H-bonding distance of E2 E531 (Fig. 3e and Table S3). CDRH2 is buried in a hydrophobic region of CD81BL formed by E2 conserved S528, W529, E531, and N532 (Fig. 3e, Fig. S3 and Table S4) and likely makes contact to the glycan at N532 (Table S3). The side chain of T54 H-bonds with E2 E531 (Fig. 3e and Table S3). R56 makes H-bond and charged interactions with E2 E531 and the main-chain carbonyl of G53 is within H-bonding distance of the N532 carbohydrate (Fig. 3e and Table S3). CDRH3 W100 is positioned adjacent to a hydrophobic cluster in E2 FL L427, C429, L438, and F442, and forms an H-bond with the main-chain carbonyl of W420 (Fig. 3e and Tables S3, S4). CDRH3 G97, S98, and W100 are involved in a network of vdW contacts with E2 FL L438 and CD81BL W529 (Fig. 3e and Table S4). The RM5-16 LC contacts E2 via all three CDRLs (Fig. 3e). CDRL1 hydrophobic residues also bind into the hydrophobic pocket formed by E2 FL L438, A439, and F442 (Fig. 3e and Table S4). In LFR3, K53 forms a H-bond with E2 H421 while CDRL3 N93 and S94 form H-bonds to E2 FL N434 (Fig. 3e and Table S3).

### Affinity of RM AR3-targeting Fabs for HCV E2

Previous studies using biolayer interferometry (BLI) to characterize the binding of RM1-73 and RM11-48 IgGs to HCV E2 showed the Abs bound strongly to all tested isolates with nM K_d_ values ^45^. To avoid avidity issues of the IgG format, in this study, we tested binding of E2 with five RM Fabs. Our BLI data show that Fabs RM1-73, RM11-48, and RM10-30 bind to E2 proteins from different isolates and genotypes including H77 (1a), HCV-1 (1a), J6 (2a), S52 (3a), and HK6a (6a) in the nM range (Fig. 2f and Table S5). Fabs RM1-73 and RM10-30 generally showed stronger binding than RM11-48, RM1-36, and RM5-16 (Fig. 2f and Table S5). Consistent with our previous study, IgG RM1-73 also exhibited enhanced binding to E2 compared to RM11-48 ^45^. Fab RM5-16 showed nM binding to all tested strains except H77 (1a) (Fig. 2f and Table S5), aligning with previous neutralization assays ^45^. Overall, we observed reduced/no binding of RM11-48, RM10-30, RM1-36, and RM5-16 to E2 from the H77 strain (Fig. 2f and Table S5), likely due to the E531A substitution in the E2 CD81BL of H77 strain (Fig. S3). The negatively charged glutamate residues at position 531 is a key part of the epitope for these Abs, except for RM1-73 (Fig. 3 and Table S3). Consequently, RM1-73 retains strong binding to the H77 strain, while the others are affected (Fig. 2f and Table S5). Sequence analysis further revealed that glutamate residue is found at position 531 in about 65% of all HCV E2 sequences extracted from the BV-BRC database (https://www.bv-brc.org/). Alanine is the next most common residue at this position, found in about 15% of sequences. These findings underscore the importance of HCV E2 residue 531 and the role of the HCV natural polymorphisms in modulating RM Ab recognition and sensitivity.

### RM AR3-targeting Abs mimic neutralizing Abs isolated from human elite neutralizers

Previously described human elite neutralizer mAbs 1198 and 1382 share similar binding modes to RM Fabs RM2-01 and RM11-43 ^20^. To further understand differences/similarities in how these five RM Abs bind to E2, we compared RM Ab-E2 structures to previously reported AR3-targeting RM and human IGHV1-69 class Abs. All five RM Abs target the same AR3 region as the previously reported RM2-01, RM11-43, as well as human bnAbs AR3C, HEPC74, mAbs 1198, and 1382 (Fig. 4a and Fig. S4) ^20,26,45,50^. The RM Abs and human elite neutralizer mAb1198 share similar overall binding poses with mAb1382 but differ in their precise approach angles relative to mAb1382 (Fig. S4). These approach angles are defined here between the center of mass of the respective Fabs and E2, ranging from counterclockwise (up to 26°) to clockwise (up to 15°) (Fig. S4). In contrast, all of these Fabs adopts markedly different approach modes compared to human bnAbs AR3C and HEPC74 (Fig. S4). When bound to E2, HEPC74 is rotated about 170° along the Y-axis relative to RM Abs, causing switching of the relative positions of HC and LC, while also exhibiting an approximately 10° clockwise rotation relative to mAb1382. Additionally, AR3C displays an ∼90° rotation along the Y-axis relative to RM Abs (Fig. S4).

**Fig. 4:**
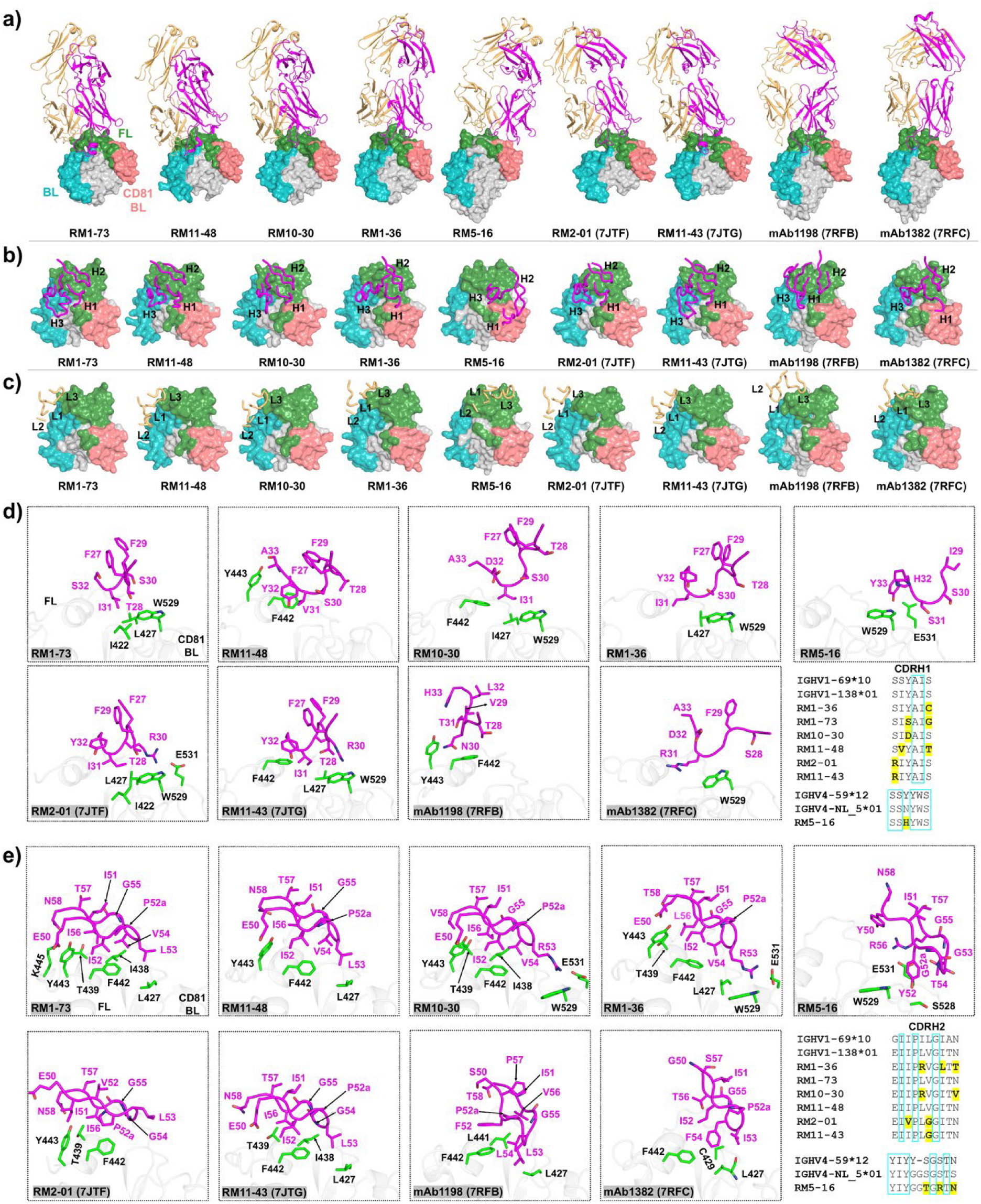
RM IGHV1-138*01 Ab recognition of E2 mimics human elite neutralizers mAbs 1198 and 1382, while IGHV4-NL_5*01 RM5-16 has a new binding mode. **a** The E2 from all complex structures were superimposed to compare the binding modes of RM Fabs and human mAbs 1198 and 1382. Fab backbones are colored with HC (magenta) and LC (light orange). E2 proteins are shown as surfaces with different colors of the subregions as in Fig. 2. **b-c** The positions of H1-3 and L1-3 of RM Abs on E2 proteins are compared with human mAbs1198 and 1382. H and L are shown in magenta and gold ribbon representations, respectively. **d-e** The binding mode of RM Ab CDRH1-2 are compared with those of human mAbs 1198 and 1382. Key E2 residues are in green sticks and CDRH1-2 are in magenta sticks. Sequences of CDRH1,2 of RM Abs are defined in Figs. 1, 2. Bottom-right panel: CDRH1-2 sequence alignment of mature RM Abs, their inferred germline counterparts, and corresponding human germline homologs. Residues conserved between RM and human germlines are boxed in cyan; somatic mutations in the mature RM Abs relative to their RM germlines are highlighted in yellow.

RM1-36 closely matches the approach angle of human mAb1382 (Fig. 4a and Fig. S4a). Similar to RM2-01 and RM11-43, RM1-73, RM11-48, and RM10-30 rotate away from mAb1382 toward the BL at counterclockwise angles of 22°, 26°, and 18°, respectively (Fig. S4a). By contrast, RM5-16 rotates away from mAb1382 towards the E2 CD81BL at a clockwise angle of 15° (Fig. S4a). As the CDRH3 dominates the binding interactions of AR3-targeting Abs (Fig. 2c), we also performed structural alignments between the CDRH3 regions of RM Abs compared to human mAb1382 (Fig. S4b). Similarly, the alignments revealed CDRH3 of RM1-36 has the same approach angle as that of mAb1382, while CDRH3s from other Abs have different binding approach angles in comparison with mAb1382 (Fig. S4b).

When we compared the position of CDRH1-3 loops of these RM IGHV1-138*01 class Abs upon interacting with E2 protein, we found their CDRH3 loops contact the BL and central β-sandwich and lower FL region, similar to the CDRH3 loops of mAbs 1198 and 1382 (Fig. 4b). In contrast, CDRH3 loops in human AR3C and HEPC74 contact the FL and CD81BL regions of E2 ^26,50^. For IGHV4-NL_5*01 RM5-16 Ab, the CDRH1-2 loops shift towards the E2 CD81BL center and lower FL region (Fig. 4b). The CDRH3 loop of RM5-16 is centrally located in a cleft between E2 FL, CD81BL, and BL (Fig. 4b). The CDRL1-3 loops of RM5-16 lie in the middle of the FL region of E2, unlike other those of other RM IGHV1-138*01 class Abs (Fig. 4c).

When we compared the CDRH1 loops of all RM AR3-targeting Abs, we observed that while RM1-73, RM2-01, and RM1-36 only contact E2 residues from the E2 FL N-terminus and CD81BL, the those of RM11-48, RM10-30, and RM11-43 extend their contacts to residues from the helix α1 C-terminus of E2 (Fig. 4d). As previously mentioned, the CDRH1 loop of RM5-16 only contacts E2 CD81BL (Fig. 4d). Meanwhile, the CDRH1 loops of RM11-48 and mAb1198 share a similar binding mode as they both contact only the helix α1 C-terminus (Fig. 4d).

Next, we compared the CDRH2 loops and found that all RM Abs have slightly different approach angles for CDRH2 (Fig. 4e). The CDRH2 loops of RM1-36 and RM10-30 bind at a similar location as those of mAbs 1382 and 1198 (Fig. 4e). In contrast, CDRH2 of RM2-01 interacts only with the E2 FL helix α1 (aa 438-443), whereas CDRH2 of RM1-73, RM11-48, RM11-43, and RM1-36 extend their footprint to E2 FL N-terminus (aa 427-429), similar in footprint but not in angle to mAbs 1198 and 1382 (Fig. 4e and Table S4). Notably, the CDRH2 tip of RM10-30 and RM1-36 also contacts E2 CD81BL W529 whereas the RM5-16 CDRH2 tip only interacts with E2 CD81BL (Fig. 4e and Table S4).

### AR3-targeting Abs compared to CD81 receptor binding

Clearly, the structures and competition data show that RM AR3-targeting Abs compete with host CD81 receptor binding to E2 ^44,45^. To further understand how these RM Abs can block CD81 receptor binding and to rationalize their relative neutralization activity, we superimposed the E2 structures from complexes with all RM Abs (RM1-73, RM11-48, RM10-30, RM1-36, RM2-01, RM11-43, and RM5-16), human AR3-targeting Abs (AR3C, HEPC74, mAbs 1198, and 1382), and CD81 receptor long extracellular loop (PDB ID 7MWX) (Fig. 5a). As expected, all RM AR3-targeting Abs target a similar E2 region as the CD81 receptor (Fig. 5a) ^22^. These and other AR3-targeting Abs bind to E2 with CD81BL in a retracted conformation, unlike its structure in complex with the CD81 receptor where it displays an extended conformation (Fig. 5a) ^22^.

**Fig. 5:**
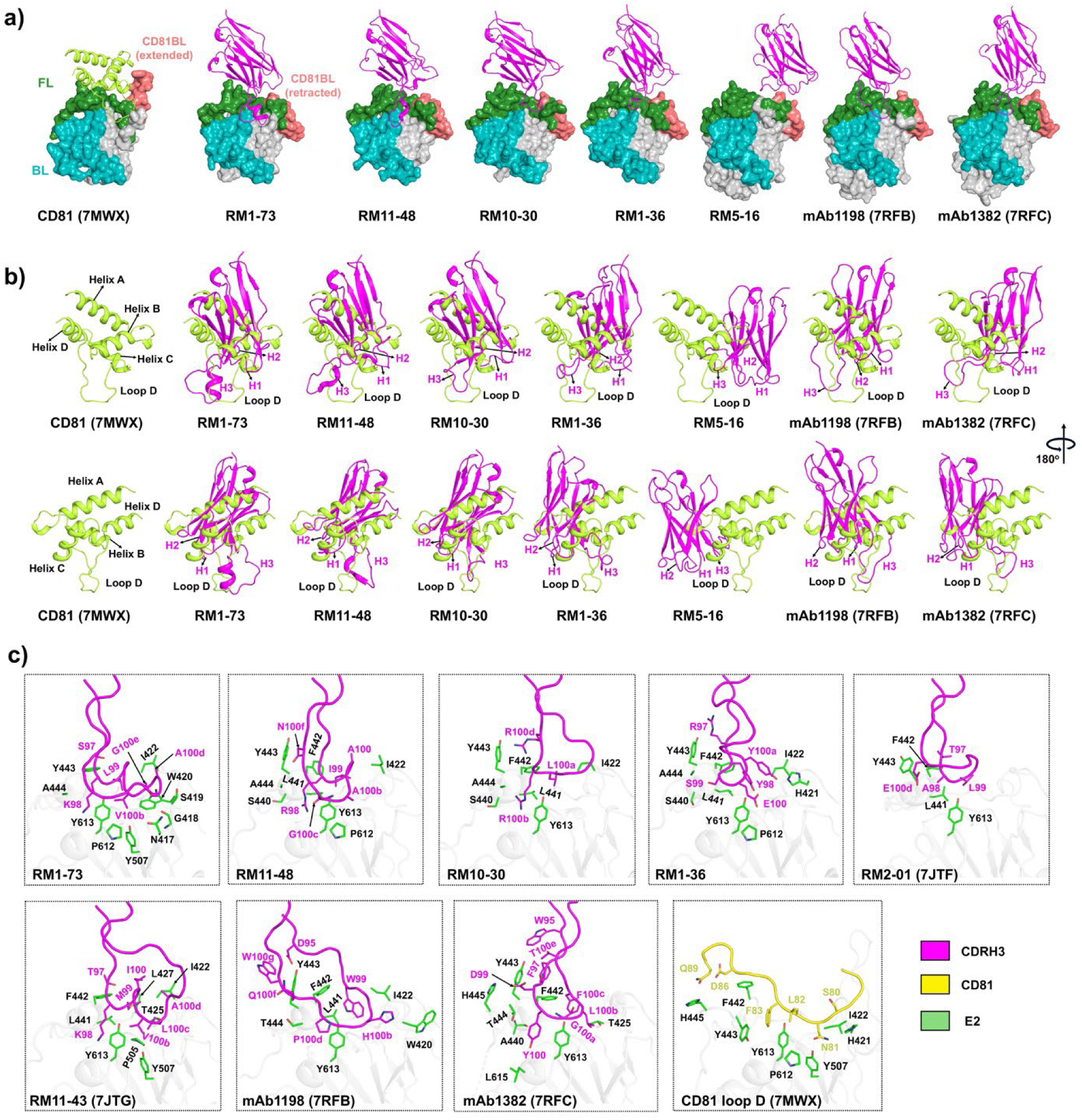
Interaction patterns of AR3-targeting Abs and CD81 receptor in complex with HCV E2. **a** The IGHVs of AR3-targeting Fab are compared with host CD81 receptor bound to E2 (PDB ID 7MWX). The CD81BL in all Fab-E2 complexes is retracted, which differs from its extended conformation in the CD81 receptor-E2 complex ^23^. The color of E2 subregions and HC and LC of Fab are as in Fig. 2. CD81 is shown in a lime cartoon representation. **b** The positions of H1-3 of AR3-targeting Abs relative to CD81. CDRH3 loops of RM IGHV1-138*01 Abs (RM1-73, RM11-48, RM10-30, RM1-36, RM2-01, and RM11-43) and human IGHV1-69 Abs (mAbs 1198 and 1382) bind to similar regions as CD81 loop D. Additionally, CDRH1-2 of these Abs interact with a region on E2 targeted by CD81 helix C. In contrast, the CDRH3 loop of RM5-16 binds near the location bound by CD81 helix C. H1-H3 and CD81 are shown in magenta and lime ribbons, respectively. **c** The binding approach and interaction of Fab CDRH3 and CD81 loop D with E2. The key interface residues of E2, Fab CDRH3, and CD81 loop D are in green, magenta, and yellow sticks, respectively. CDRH3 and CD81 loop D are shown as backbone ribbons with stick side chains. E2 proteins are shown as light grey cartoons.

Loop D of CD81 binds into the hydrophobic groove of E2, which is comprised of the FL N-terminus (aa 421-422), helix α1 C-terminus (aa 442-445), β-sandwich Y507, and BL (aa 612-613) ^22^. We found that the CDRH3 loops of RM IGHV1-138*01 Abs (RM1-73, RM11-48, RM10-30, RM1-36, RM2-01, and RM11-43) and human IGHV1-69 mAbs 1198 and 1382 also bind into this groove (Fig. 5b, 5c). As a result, residues from CDRH3 of RM IGHV1-138*01 Abs, human IGHV1-69 Abs (mAbs 1198 and 1382), and CD81 loop D form somewhat similar interactions with E2 (Fig. 5c and Table S4) ^22^. The CD81 helix C interacts with E2 at the hydrophobic pocket formed by FL N-terminus (aa 426-429), FL helix α1 (aa 438-443), and CD81BL (aa 523-531) ^22^ and the CDRH2 regions of IGHV1-138*01 Abs (RM10-30 and RM1-36) make a similar interaction (Figs. 4e, 5b).

### Similarities and differences within AR3-targeting Abs

Structural analyses of the RM Ab-E2 complexes show similar Fab approach angles and epitope conservation. Although IGHV4-NL_5*01 encoded RM5-16 uses a different HC germline gene, it recognizes a similar but not identical epitope as RM IGHV1-138*01 Abs (Fig. 3). For example, RM5-16 shares some common epitope residues such as FL L427, I438, A439 (A439 in HCV-1 strain), F442, Y443, and CD81BL W529 (Fig. 3 and Table S4). RM5-16 also shares numerous common epitopes within the FL loop (aa 429-438) with human AR3-targeting IGHV1-69 Abs such as AR3C, HEPC74, mAbs 1198, and 1382 (Fig. 3e) ^16,20,26^. Interestingly, the CDRH3 tip of RM5-16 inserts into E2ecto at a location similar to that bound by AR3C and HEPC74 CDRH3s (Fig. 3e and Table S4) ^16,26^.

Six RM IGHV1-138*01 Abs (RM11-48, RM1-73, RM1-36, RM10-30, RM2-01, RM11-43) target overlapping conserved epitopes including E2 FL (I422, T425, L427, I438, T439, L441, F442, Y443, and A444); E2 β-sandwich (P505), CD81BL (W529), BL(Y613) (Fig. 3, Fig. S3, and Table S4) ^20,45^. The conserved epitope residues E2 FL A/S440 (A440 in E2c3 HK6a; S440 in E2ecto 1b09 strain) and BL (P612) are present for RM1-73, RM11-48, RM1-36, RM11-43, mAbs 1198, and 1382, but not RM2-01 (Fig. 3, Fig. S3, and Table S4) ^20,45^.

RM1-73 has the longest CDRH3 (25 aa) among RM AR3-targeting nAbs studied here (Fig. 1f). In the complex of RM1-73 with HK6a E2c3, we found ordered electron density for E2 AS412 (aa 417-420), β-sandwich loop (aa 542-550), and post variable region 3 (pVR3) (aa 566-576), that are disordered in other RM IGHV1-138*01 Fab-HK6a E2c3 complex structures (Fig. S5a). In previous crystal structures, the AS412 region has displayed many conformations (e.g. beta-hairpin, semi-open, and open) ^51^. In the low pH structure of CD81-E2, one complex in the asymmetric unit shows AS412 residues 415-424 folding around the CD81 loop D, facilitating interaction of E2 with the CD81 receptor (Fig. S5b) ^23^. The second CD81-E2 complex in the asymmetric unit has density for E2 starting at residue 418, with residues 418-424 forming the same interaction with CD81 ^23^. Upon RM1-73 binding, AS412 residues 417-424 adopt an extended conformation, following the general path of AS412 in the CD81 complex, binding close to the tip of CDRH3, bringing E2 N417, G418, S419, and W420 into close contact with RM1-73 (Fig. S5b and Table S4). The CD81 loop D and the long CDRH3 of RM1-73 may serve to stabilize the flexible AS412 region (Fig. S5a and S5b), at least in the crystal structures.

Although not in the Fab epitope, the HCV-1 E2ecto AS412 region, as well as the VR2 and VR3 loops have ordered electron density in the RM5-16 complex structure (Fig. S5c). These regions are disordered in structures in complex with human mAbs 1198 and 1382 (Fig. S5c).

While the HCV E2 FL has been seen to adopt an alternate ‘B’ conformation in complex with human bnAbs 212.1.1 and HC1AM ^21^, it is found in the more common ‘A’ conformation in our seven RM Ab structures targeting AR3 as well as in the CD81 receptor complex structure (Fig. S6a).

### The motifs and somatic mutations in HCs of AR3-targeting RM Abs which are critical for E2 binding and virus

Both germline genes of AR3-targeting RM Abs have hydrophobic residues in their CDRH2: IGHV1-138*01 encodes I^51^I^52^P^52a^L/R^53^V^54^G^55^I^56^T^57^ and IGHV4-NL_5*01 encodes I^51^Y^52^G^52a^G^53^S^54^G^55^S^56^T^57^ (Figs. 1f, 1g, and 4e). It has been noted that the hydrophobic CDRH2 tip of human IGHV1-69 AR3 and AS434-targeting Abs plays an important role for recognizing conserved hydrophobic epitopes on E2 ^25^. Similar to human IGHV1-69 bnAbs, most RM IGHV1-138*01 AR3-targeting Abs have hydrophobic CDRH2 loops that pack into a hydrophobic pocket surrounded by the FL and CD81BL (Fig. 4e and Fig. S7a). In RM10-30 and RM1-36 Abs, CDRH2 L53 undergoes somatic mutation to R53 (Fig. 4e and Fig. S7a) enabling RM10-30 and RM1-36 to form a salt bridge with E531 and hydrophobic interaction with W529 of E2 CD81BL (Figs. 3, 4e and Fig. S7a). A V54G somatic mutation occurs in RM2-01 and RM11-43 removing the hydrophobic interaction of V54 with E2 FL (Fig. 4e and Fig. S7a).

CDRH1 S30 (in RM1-36, RM1-73, RM10-30, and RM11-48) is somatically mutated to R30 in RM2-10 and RM11-43 (Fig. 4d and Fig. S7a). This R30 side chain occupies a similar spatial position as CDRH2 L/R53 of RM1-73, RM11-48, RM10-30, and RM1-36 (Fig. S7a). All of these Arg residues make similar interactions with the FL and CD81BL regions (Fig. S7a) ^45^.

In CDRH3 of AR3-targeting RM Abs, the basic, long side chains of K98 (RM1-73), R100b (RM10-30), R98 (RM11-48), and K98 (RM11-43) bind to similar regions in E2 (Fig. 5c and Fig. S7b). In contrast, the β and γ atoms of the small S99 (RM1-36) and A98 (RM2-01) are found in similar positions as the longer side chains K98/R98/R100b (Fig. 5c and Fig. S7b). Notably, these residues engage in extensive interactions with the E2 hydrophobic pocket formed by FL (L441, F442, Y443, and A444), which is also the binding site for the aromatic F83 of CD81 (Fig. 5c). Mutating CDRH3 residues R98A (in RM11-48) and K98A (in RM11-43) substantially reduced both the stability (or melting temperature (Tm)) and binding affinity to E2 (Fig. S7d, S7e). While the RM10-30 R100bA mutation did not affect the Tm, its binding affinity to E2 decreased nine-fold compared to the RM10-30 wild-type (WT) (Fig. S7d, S7e). The RM1-73 K98A mutant retained Tm and binding affinity to E2 compared to RM1-73 WT (Fig. S7d, S7e), likely due to main chain H-bond interactions from RM1-73 K98 to E2 (Fig. 3a and Table S3). In contrast, RM1-36 S99R and RM2-01 A98R mutants did not have much effect on either Tm or binding affinity to E2 when compared to the WT Fabs (Fig. S7d, S7e). Previous studies revealed that mutation of K98R in RM11-43 substantially improved its ability to bind to and neutralize heterologous viruses ^45^, suggesting that the positive long side-chain CDRH3 K/R of RM Abs play a key role in E2 binding and virus-neutralization.

The RM5-16 CDRH2 SHM S54T and S56R play an important role in E2 binding (Fig. 3e and Tables S3, S4). For example, CDRH2 T54 forms H-bonds and vdW contacts with CD81BL E531 and N532 while R56 is involved in extensive vdW and salt bridge interactions with E531 (Fig. 3e and Tables S3, S4). R56 of RM5-16 binds to CD81BL E531 in a similar manner as CDRH2 R53 of RM10-30, RM1-36, and CDRH1 R30 in RM2-01 and RM11-43 (Fig. 3 and Table S4). We also observed that RM5-16 CDRH3 W100 is positioned in a similar location as the key CDRH3 disulfide motif in human IGHV1-69 class Abs AR3C and HEPC74 (Fig. S7c) ^16,26^, close to a pocket formed by E2 aromatic residues (L427, L438, L441, and F442) and the E2 C429-C503 disulfide bond. The CDRH3 W100A mutation had no affinity for E2, but its Tm relative to RM5-16 WT was not affected (Fig. S7d, S7e). This result supports the idea that RM5-16 CDRH3 W100 is critical for its role in E2 binding, potentially similar to the disulfide motif of human IGHV1-69 AR3C and HEPC74 (Fig. S7c) ^16,26^.

Collectively, our sequence and mutation analyses indicate that specific somatically derived amino acids in RM AR3-targeting nAbs that are positioned at the Ab-E2 binding interface play important roles in mediating E2 binding as well as in neutralization activity.

## Discussion

One of the challenges for HCV vaccine research is the absence of an immunocompetent animal model for preclinical immunization and protection studies ^4,52,53^. The extensive genetic diversity of HCV, along with the flexibility of the E2 glycoprotein, may necessitate rational design of a cross-genotype vaccine that targets conserved epitopes ^4,21,53,54^. The rational design and evaluation of such HCV vaccine immunogens can be guided by structural information of nAb epitopes ^18,25,50,53,55–58^.

Compiling information about shared germline gene segments, structural features, and antigen-interaction patterns of nAbs derived from humans or animals is important for guiding effective HCV immunogen design. These data are increasing as more human bnAbs against HCV are identified from both chronically and spontaneously infected HCV patients ^14,25–27^. Anti-HCV human or human-like IGHV1-69 class Abs, from both phage display and HCV-infected patients or NHPs, seem to preferentially target the conserved AR3 region of E2 ^25,44,45^. Human IGHV1-69 AR3-targeting Abs also bind to a similar epitope with most of the contacts from their HCs ^25^. Human IGHV1-69 AR3-targeting nAbs possess signature hydrophobic sequences consisting of CDRH2 I/V/T^52^-P^52a–53^-F/S^54^ (X = hydrophobic residue) and an aromatic residue Tyrosine (Y) in CDRH3 that insert into the hydrophobic AR3 of E2 ^25^. Recently, structures for FL-binding human Abs hcab55 and hcab64 encoded by the human IGHV1-46 germline gene have been reported ^27^. Although encoded by a different IGHV gene, IGHV1-46 class nAbs utilize similar approach angles to recognize overlapping epitopes in the E2 FL as human IGHV1-69 class bnAbs HEPC74 and AR3C ^27^.

Our RM Ab structural analyses reveal that, like human IGHV1-69 class Abs, the RM IGHV1-138*01 class Abs primarily use their HC to interact with E2 (Fig. 2c). CDRH3 contributes substantial BSA in most RM IGHV1-138*01 (or human IGHV1-69) class Abs (Fig. 2c). Importantly, the RM IGHV1-138*01 is highly similar to the human IGHV1-69 germline gene, and their mature Abs have very low SMH (∼1-4 %) (Fig. S1d). Basic K/R and hydrophobic residues in CDRH3 and a hydrophobic motif I/V^52^P^52a^L/R^53^V/G^54^ at the CDRH2 tip interacts with the conserved and hydrophobic AR3 region of E2. Their long K/R CDRH3 residues can reach into and extensively interact with E2 in the AR3 hydrophobic groove through multiple types of interaction such as H-bonds, vdW, and hydrophobic interactions (Fig. S7b), facilitating the specificity and affinity of the antigen-Ab interactions. Some key CDRH1 and CDRH2 residues have exchangeable roles on binding E2 (Fig. S7a).

Notably, these RM IGHV1-138*01 class Abs share very similar binding modes and epitopes with human IGHV1-69 Abs from human elite neutralizers ^20^. Although NHP-derived AR3-targeting Abs approach the E2 neutralizing face with slightly different orientations and angles, they recognize a shared group of conserved AR3 residues with human anti-HCV IGHV1-69 bnAbs. As NHPs share many similarities with humans in their immune systems ^32^, the functional and structural similarities of the antibodies described here suggest that the NHP model can to some extent recapitulate the human immune response to the same HCV antigen and is a valuable preclinical study model to study how Abs are elicited by HCV vaccine candidates.

Anti-HCV AR3-targeting Abs were previously grouped into two classes based on their CDRH3 loop shape and secondary structure: either straight β-hairpin (HEPC3, HEPC74, hcab55, hcab64, and AT1209) ^5,26,27^ or bent β-hairpin (AR3A, AR3C, and HC11) ^16,21,24^. CDRH3s for RM1-73, RM11-48, and RM11-43 incorporate helical segments when bound to E2 (Fig. 4a and Fig. S2a) ^45^, in contrast to loop conformation of the unliganded RM1-73 Fab (Fig. S2a). Like CDRH3 loops of mAbs 1198 and 1382, those of RM10-30, RM2-01, RM1-36, and RM5-16 have loop conformations when bound to E2 (Fig. 5a). These observations suggest that the shape and length of the CDRH3 loop may influence the approach angles of Abs to the antigen.

RM5-16 from a different immunized animal arose from the IGHV4-NL_5*01 germline (a homolog of human IGHV4-59*12) but targets the same AR3 region using interactions from both HC and LC (Figs. 2e, 3e). As noted, IGHV4-NL_5*01 germline alleles appear 3.4-fold more frequently than IGHV1-138*01 germline alleles in the animals contributing to the MUSA dataset (Fig. 1d). Moreover, the human IGHV4-59 germline gene was also found to be the most frequently used among B cell receptors in chronic HCV patients, both prior to and during DAA treatment ^59^, indicating that RM IGHV4-NL_5*01 or human IGHV4-59 germline genes may be prevalent in effective Ab-mediated responses against HCV. To date, structural information for anti-HCV human IGHV4-59 as well as NHP IGHV4-NL_5*01 class Abs remains sparse. Noticeably, unlike IGHV1-138*01 Abs, RM5-16 does not interact with the E2 BL C-terminal region (aa 612-616), but instead makes extensive contact with the CD81BL (Fig. 3e). E2 CD81BL Y529 and W531 are thought to be proximal to the membrane and involved in membrane binding ^22,23^, suggesting that RM5-16 might have an alternative neutralization mechanism beyond most characterized AR3-targeting class Abs. Of note, despite different angles of approach, we found that the IGHV4-NL_5*01 (a homolog of human IGHV4-59*12) and IGHV1-138 *01 (a homolog of human IGHV1-69*10) class Abs have similar binding footprints and occupy the same set of conserved hydrophobic pockets in the AR3 region of E2, confirming convergent recognition in NHPs as well as humans. Notwithstanding, the binding mode of the NHP IGHV4-NL_5*01 class Abs might be specific to NHPs or not seen yet in humans. Thus, RM5-16 offers another starting point for uncovering analogous human Ab responses to HCV infection.

Importantly, all RM AR3-targeting Abs studied here block CD81 binding to E2 ^44^ and exhibit a binding mode analogous to that of the host CD81 receptor (Fig. 5). Collectively, host receptor functional binding mimicry of AR3-targeting Abs highlights parallels between receptor engagement and Ab neutralization.

While HCV does not naturally infect RMs, limiting their use for studying the full viral life cycle, our structural analysis of RM Ab-antigen complexes, alongside comparisons with human Abs and host receptor, provide valuable insights into the immune response to HCV immunogens. Our findings offer critical preclinical data and provide a baseline for evaluating the ability of vaccine candidates to elicit nAbs and protect against diverse HCV strains.

## Methods

### IG-targeted capture genome sequencing, assembly, and analysis protocol

From the RM PBMCs, 2.5 ug of high molecular weight DNA was sheared using a g-tube (Covaris, Woburn, MA, United States) to ∼14 Kbp at 4000 RPM and size selected via bead-based size selection to remove fragments less than 3 Kbp. Sheared gDNA under-went end-repair and A-tailing using the standard KAPA library protocol (KAPA Hyper Prep Kit; Roche, Indianapolis, IN, United States). Barcodes were added to samples sequenced on the Sequel IIe platform and universal primers ligated to all samples. PCR amplification was performed for 8 cycles using PrimeSTAR GXL Polymerase (Takara, San Jose, CA, United States) at an annealing temperature of 60 °C. Small fragments and excess reagents were then removed using 0.7X vol: vol KAPA Pure beads (Roche). Genomic DNA target enrichment was carried out using oligo probes designed directly from the reference sequence for IGH/K/L of rhemac10 (53) (IGH (chr7:167,854,585-169,917,761), IGK (chr13:16,744,193-18,180,859), IGL (chr10:29,581,424-30,956,134), and assembly following Cirelli et al. (54). Constructed capture libraries were washed using the KAPA Hyper-Cap protocol (KAPA HyperCapture Reagent kit and KAPA HyperCapture Bead kit, Roche), and post-capture PCR amplification performed for 15 cycles using PrimeSTAR GXL Polymerase (Takara) at an annealing temperature of 62 °C. Sequencing SMRTbell libraries were prepared using the SMRTbell Template Prep Kit 2.0 and SMRTbell Enzyme Cleanup Kit 1.0 (Pacific Biosciences, Menlo Park, CA, United States). Each sample was treated with a DNA Damage Repair and End Repair mix to repair nicked DNA, followed by A-tailing and ligation with SMRTbell hairpin adapters. These libraries were treated with an exonuclease cocktail to remove unligated gDNA and cleaned with 0.6X AMPure PB beads (Pacific Biosciences). The resulting SMRTbell libraries were prepared for sequencing according to the manufacturer’s protocol and sequenced on the Sequel IIe system using 2.0 chemistry and 30 hr movies. HiFi data, consisting of circular consensus sequences filtered at a quality threshold of QV20 (99%), were generated on the instrument and used for all downstream analysis.

HiFi sequencing reads were assembled into haplotype-resolved assemblies using Hifiasm ^60^ with default parameters. The IG loci were extracted and annotated using Digger ^49^. Sequences of Abs for each animal were aligned to their respective IG assemblies using BLAT ^61^. The locations of the highest BLAT matches were linked to the corresponding germline allele Digger annotations. Allele assignments were based on names used by KIMDB ^38^ and MUSA ^48^ databases, both of which included exact matches to the closest germline identified in the assemblies of each animal. The SHM for the RM Abs were assigned by IgBLAST with the input of *Macaca mulatta* (IGHV, D, and J germline gene) sequence files from the sequenced germline alleles. All HC and LC sequences of RM Ab in this study are aligned by Clustal Omega ^62^. The Fab residues were renumbered according to Kabat nomenclature ^63^.

### Protein expression and purification

RM Fab-derived V_L_ and V_H_ DNA fragments were inserted into the pHCMV mammalian cell expression vector containing the corresponding human λ or κ CL region and the human IgG C_H_1 region, terminating with a stop codon following the CH1 sequence ^44,64^. To enhance crystallization, we introduced the ‘CK’ mutation into κ CL regions of RM1-73, RM10-30, RM11-48, RM1-36, and HEPC46 Fabs ^65^. The HC and LC encoding plasmids for each Fab were co-transfected into Expi293F cells using ExpiFectamine (Thermo Fisher Scientific) according to the manufacturer’s instructions. The cells were grown and harvested after 7 days. Affinity chromatography using CaptureSelect CH1-XL affinity matrix (Thermo Fisher Scientific) was used to purify recombinant Fabs from culture supernatant, followed by size-exclusion chromatography (SEC) using a Superdex 200 column (GE Healthcare).

The HK6a (genotype 6a) E2c3 proteins were purified as previously described ^24^. E2 proteins from isolates H77 (1a), HCV-1 (1a), J6 (2a), and S52 (3a) were prepared for biophysical assays as reported for the HK6a E2c3 protein. For HCV-1 E2ecto and HEPC46 Fab, the complex was expressed by co-transfecting expression vectors encoding HEPC46 Fab and untagged E2ecto (residues 384-643) from strain HCV-1. HEPC46-E2ecto complexes were then purified from Expi293F cell supernatants using CaptureSelect CH1-XL affinity matrix followed by SEC. All purified proteins were quantified by optical absorbance at 280 nm and stored in 20 mM Tris pH 8.0 and 150 mM NaCl (TBS) at −80 °C for further experiments. The purities of recombinant proteins were analyzed by reducing and nonreducing SDS-PAGE.

### Biolayer interferometry

E2 binding with Fabs was evaluated by BLI using an Octet Red instrument (ForteBio) at 10 °C ^45^. The Fabs at 20 μg/mL in 1× kinetics buffer (1× PBS, pH 7.4, 0.01% BSA, and 0.002% Tween 20) were immobilized onto Fab-2G biosensors and interacted with a two-fold gradient dilution of E2 analyte proteins starting at 540 nM. The kinetic assay consisted of the sequential steps: baseline equilibration (60 s, buffer), Fab protein immobilization (180 s), washing unbound Fab proteins (30 s, buffer), second baseline stabilization (60 s, buffer), E2 binding association (600 s), and E2 dissociation measurement (600 s, buffer). For estimating the dissociation constant (K_d_), a 1:1 binding model was used.

### Differential scanning fluorimetry (nanoDSF)

A Promethius Panta (nanoTemper Technologies) was used to measure the melting temperatures of the wild-type (WT) and mutant RM Fabs. The WT and mutant Fabs at 1 mg/ml in TBS were loaded into capillaries and inserted into the sample holder. We applied a temperature gradient of 1°C/min from 25 to 95 °C for each sample. The intrinsic protein fluorescence at 330 and 350 nm was recorded. Three independent measurements were performed for each sample. The data was analyzed by Panta Analysis v1.8. The melting point (Tm) was determined as the maximum of the first derivative of the melting curve.

### Crystal structure determination

For complexes of RM1-36, RM10-30, RM11-48, and RM1-73 with HK6a E2c3, E2 proteins were combined with Fab in a 1:1.2 molar ratio. For RM5-16, the HCV-1 E2ecto complex with HEPC46 was mixed with RM5-16 Fab in a 1:1.2 molar ratio. The mixture was incubated overnight at 4°C before further purification by SEC (Superdex 200 GL column) in TBS buffer to remove uncomplexed Fab. Crystallization experiments were set up using the sitting drop vapor diffusion method at 20°C. Initial crystallization conditions were obtained from trials using our robotic, high-throughput Rigaku CrystalMation system. Crystals were manually optimized using the sitting-drop vapor diffusion method at 20 °C using Cryschem plates (Hampton Research Corp). Crystallization precipitant conditions are listed in Table S6. The protein crystals were flash-cooled at 100 K with 20% (w/v) ethylene glycol added as cryoprotectant.

Diffraction data were collected at synchrotron beamlines and integrated and scaled with HKL-2000 ^66^, or for the unliganded RM1-73 Fab were collected in-house with a Rigaku MicroMax-007 generator at 1.5418 Å wavelength and a Mar345dtb area detector. Data collection statistics are summarized in Tables S1 and S2. Crystal structures were phased by the molecular replacement (MR) method using the program Phaser ^67^. The initial input models for RM Fabs were generated by Repertoire Builder (https://sysimm.ifrec.osaka-u.ac.jp/rep_builder/) ^68^. The MR template for E2 was derived from HK6a E2c3 (PDB ID: 7JTG) or HCV-1 E2ecto (PDB ID: 6MEJ). Refinement was carried out in Phenix ^69^. Structure models were examined and modified with the program Coot to assess model fit and any potential clashes ^70^. Final refinement statistics are summarized in Tables S1 and S2 and their quality analyzed with MolProbity ^71^.

### Structural analysis

Epitope and paratope residues, molecular interactions, BSA of E2, Fabs, and CD81, were identified and calculated with the Protein Interfaces, Surfaces and Assemblies (PISA) web server at the European Bioinformatics Institute (www.ebi.ac.uk/pdbe/pisa/) ^72^ followed by manual inspection. Structure Figures were created by MacPyMol (DeLano Scientific LLC). RMSD calculations were done in PyMOL following pairwise Cα alignments without excluding outliers.

## Data availability

The X-ray coordinates and structure factors have been deposited in the Research Collaboratory for Structural Bioinformatics (RCSB) Protein Data Bank under accession codes 9MS9, 9MRZ, 9MNS, 9MNU, 9MNT, 9MNQ, and 9MSC.

## Abbreviations

HCV: Hepatitis C virus
NHP: non-human primates
IgG: immunoglobulin
RM: rhesus macaques
bnAbs: broadly neutralizing antibodies
DAA: direct-acting antiviral drug
IGS: individualized germline set
MSA: multiple sequence alignment
HC: antibody heavy-chain
LC: antibody light-chain
FL: front layer
BL: back layer
CD81BL: CD81 binding loop
H-bond: hydrogen-bonds
vdW: van der Waals
SHM: somatic hypermutation
BSA: buried surface area
RMSD: root-mean-square deviation
IGHV: variable domains of antibody heavy-chain
AR3: antigenic region 3
CDR: complementarity determining region

## Acknowledgments

We thank Henry Tien for automated robotic crystal screening at The Scripps Research Institute and Xiaoping Dai for help with in-house data collection. X-ray diffraction data were collected at the Advanced Photon Source (APS) beamline 23ID-B (GM/CA CAT) and the Stanford Synchrotron Radiation Lightsource (SSRL) beamline 12-2. GM/CA CAT is funded in whole or in part with federal funds from the National Cancer Institute (Y1-CO-1020) and the National Institute of General Medical Sciences (NIGMS) (Y1-GM-1104). Use of the APS was supported by the US Department of Energy (DOE), Basic Energy Sciences, Office of Science, under Contract DE-AC02-06CH11357. The SSRL is a Directorate of SLAC National Accelerator Laboratory, and an Office of Science User Facility operated for the US DOE of Science by Stanford University. The SSRL Structural Molecular Biology Program is supported by the DOE Office of Biological and Environmental Research, and by the NIH, NIGMS (including P41GM103393), and the National Center for Research Resources (NCRR) (P41RR001209). The contents of this publication are solely the responsibility of the authors and do not necessarily represent the official views of NIAID, NIGMS, NCRR, or NIH.

This investigation used resources that were supported by the Southwest National Primate Research Center grant P51 OD011133 from the Office of Research Infrastructure Programs, National Institutes of Health. Research reported in this publication was supported by the Office of the Director, National Institutes of Health under Award Numbers S10OD028732 and S10OD032443. The content is solely the responsibility of the authors and does not necessarily represent the official views of the National Institutes of Health.

## Author contributions

Conceptualization: YTKN, FC, RLS, ML, IAW

Methodology: YTKN, FC, RLS, ML, IAW

Investigation: YTKN, FC, EG, SS, NT, LAU, RLS, CC

Visualization: YTKN, FC, RLS, CW

Supervision: RLS, CTW, ML, IAW

Writing—original draft: YTKN, FC, RLS, SS, CTW, ML, IAW

Writing—review & editing: YTKN, FC, RLS, SS, CC, CTW, ML, IAW

## Supplementary Information

**Fig S1.**
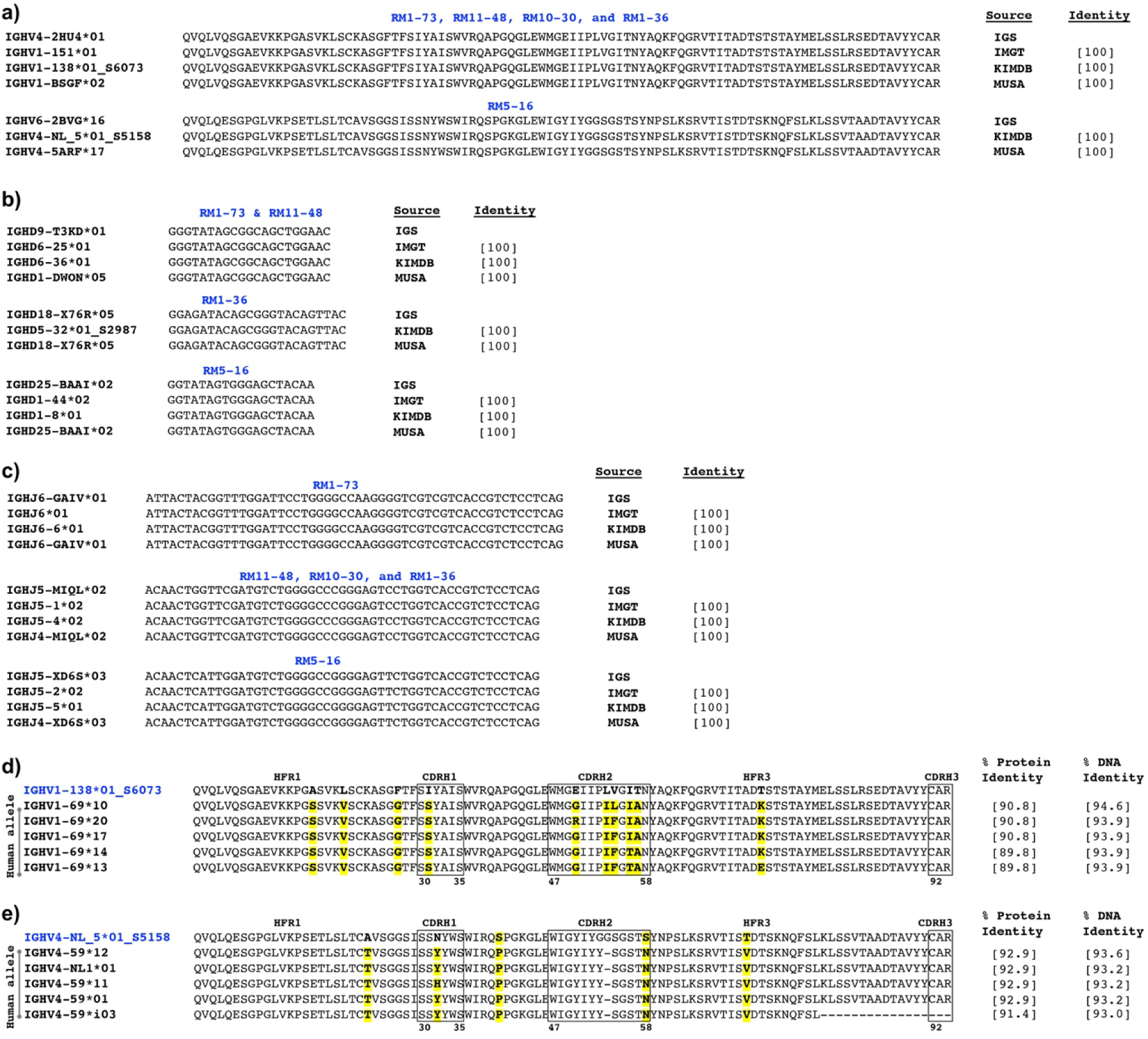
VDJ gene segment analysis of RM germline alleles and identification of closest human IGHV germline alleles, related to Fig. 1. **a-c** DNA sequence alignment of IGHV, D, J germline alleles of RM Abs derived from different sources. DNA identity comparing each IGHV, D, J germline allele to its counterpart in IGS is shown. **d**, **e** Multiple sequencing alignment (MSA) identifies five of the closest human IGHV germline alleles to RM IGHV1-138*01 and IGHV4-NL_5*01. HC sequence alignment compares the RM IGHV1-138*01 and IGHV4-NL_5*01 germline alleles with their top five closest human homologs (including IGHV1-69*10 and IGHV4-59*12 germlines, respectively). Residues that differ between RM and human germline alleles are highlighted in bold and yellow. Protein and DNA sequence identities of human and RM germline alleles are also provided. CDRH1-2 length and sequence are defined according to Kabat numbering.

**Fig S2.**
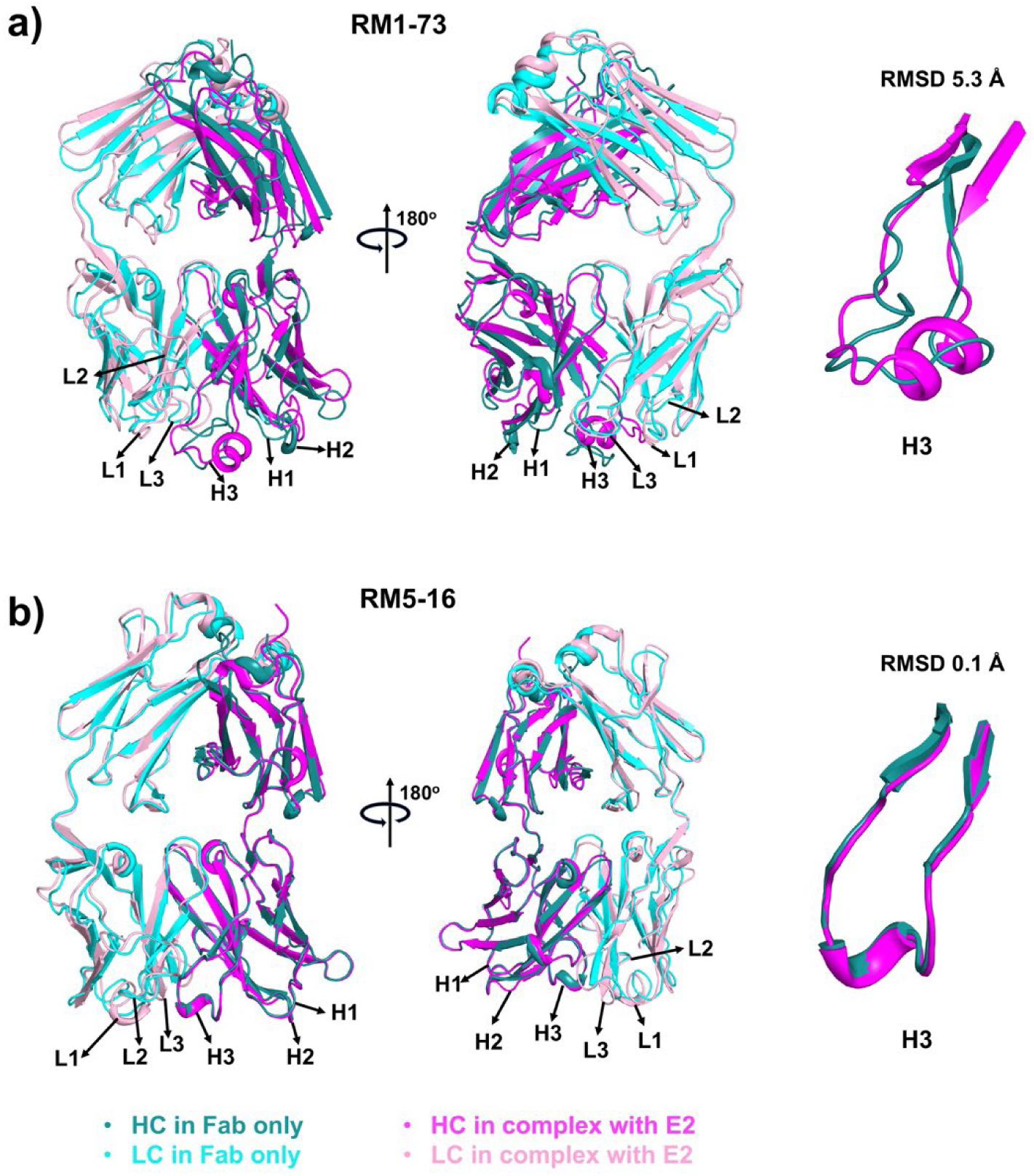
Comparison of apo and ligand-bound Fabs RM1-73 and RM5-16. **a**, **b** HC and LC in apo Fab are in teal and cyan, respectively. HC and LC of Fab in complex with E2 are in light magenta and pink, respectively. CDR H1-H3 and L1-L3 are labeled. The comparison of CDRH3 free and bound is shown on the right with Cα root-mean-square deviate (RMSD).

**Fig S3.**
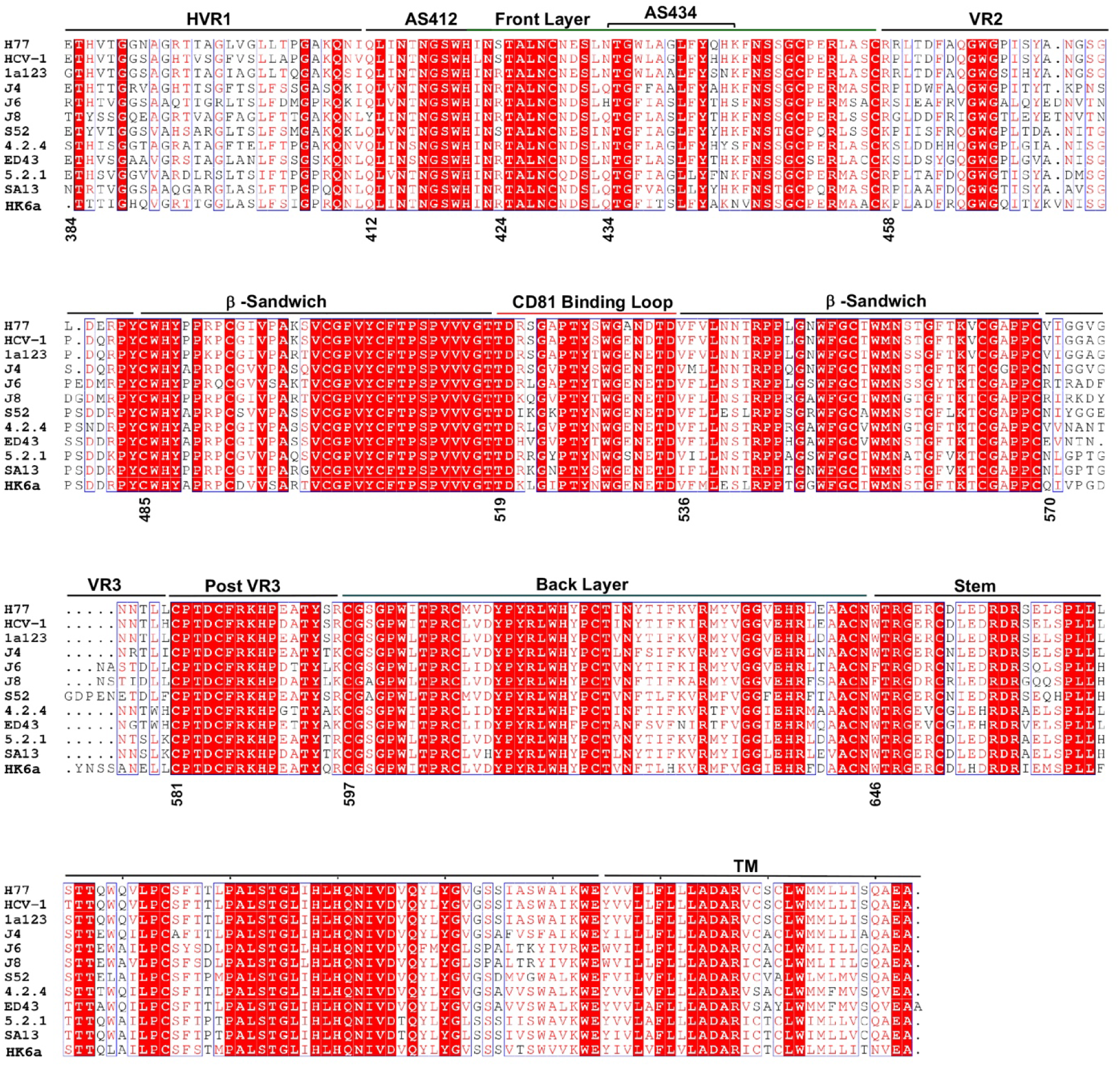
Sequence alignment of E2 across HCV isolates and genotypes. Identical residues across groups are shown as white characters on a red background. Similar residues defined by the ESPript ^73^ according to physicochemical feature are shown in blue boxes with red characters. Residues that differ at particular positions are in black characters. The E2 sequences from H77 (1a), HCV-1 (1a), 1a123 (1a), J4 (1b), J6 (2a), J8 (2b), S52 (3a), 4.2.4 (4a), ED43 (4a), 5.2.1 (5a), SA13 (5a), andHK6a (6a) were used to align. The subregions of E2 are identified and labeled as in Fig. 2a.

**Fig S4.**
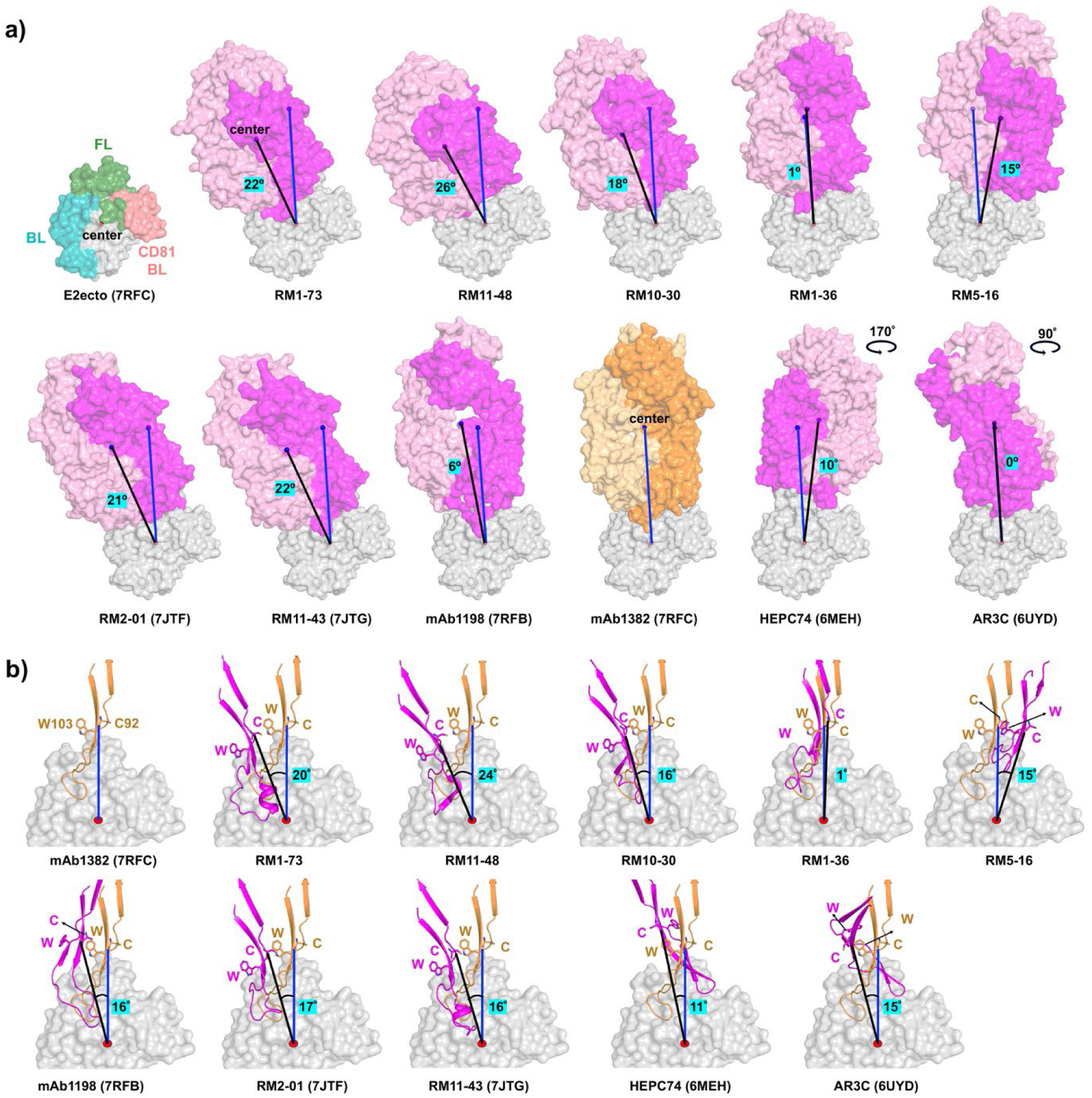
Comparison of binding approach of RM vs. human AR3-targeting Abs. **a** The RM Fabs show a similar binding angle to human elite neutralizers mAbs 1198 and 1382. The superposition of the RM Abs and human mAb1382 on HCV E2ecto protein (PDB ID 7RFC) reveal that RM Abs have similar binding angles and binding modes with mAb1382 but differ from human HEPC74 and AR3C. E2 (grey) and Fabs are shown in surface representation. HC and LC of mAb1382 are in orange and bright orange, respectively. HC and LC of all other Fabs are in magenta and pink, respectively. **b** The angle of approach of CDRH3 of all Abs on E2 is compared. Superposition of all CDRH3-containing regions (aa 87-113) with mAb1382 show different binding angles and local conformational changes of CDRH3, resulting from the differences in CDRH3 length and origin. CDRH3 is shown as a cartoon. C92 and W103 of CDRH3 in all Fabs are chosen to define the angle of approach. E2 is shown in grey surface representation.

**Fig S5.**
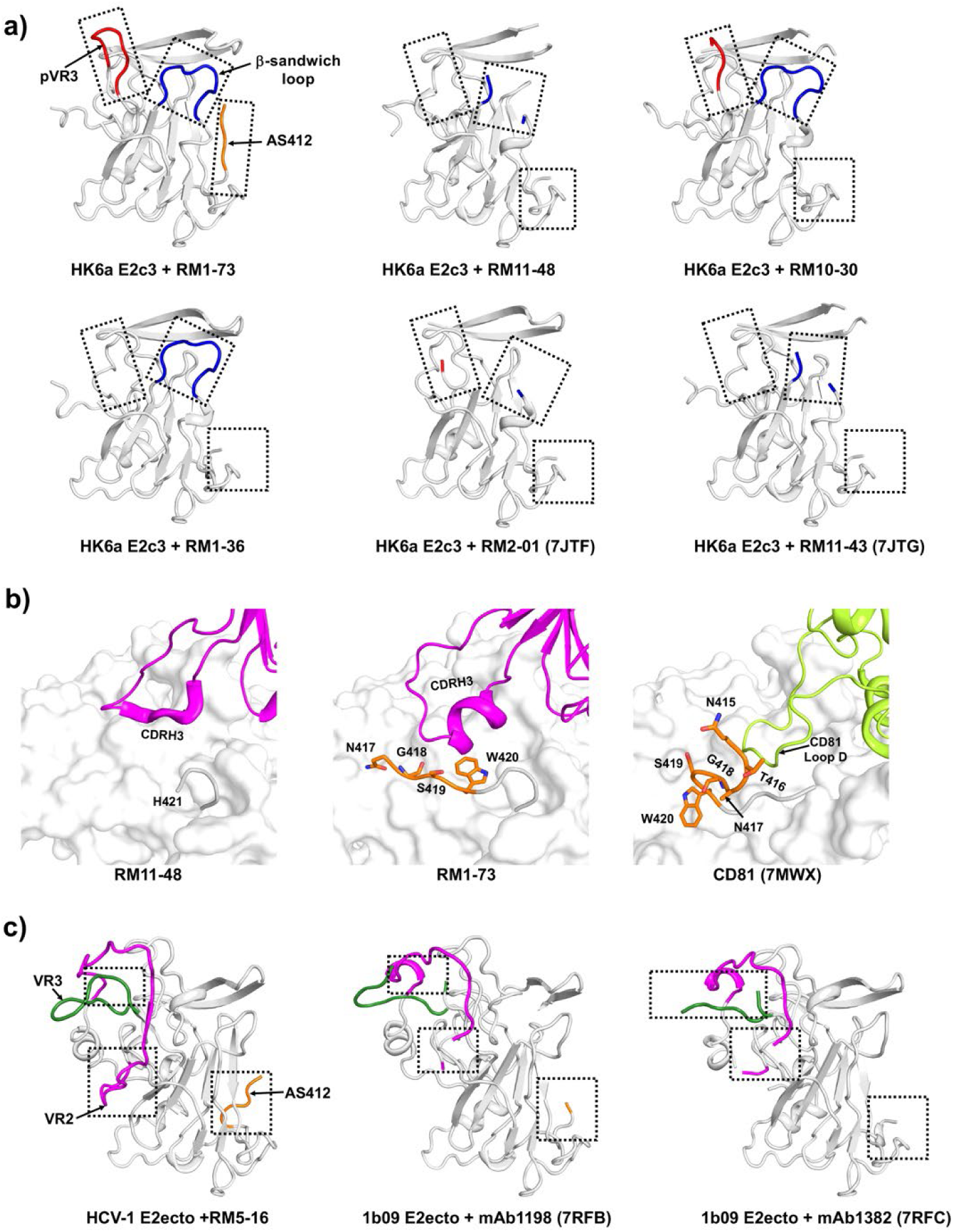
Structural alignment of E2 protein complexes with Abs and CD81 receptor shows the dynamic nature of AS412 and VR loops. **a** Alignment of HK6a E2c3 structures from complexes with different RM Abs. The AS412 (aa 412-420), β-sandwich loop (aa 542-550), and pVR3 regions of HK6a E2c3 are shown in orange, blue, and red cartoons, respectively. All remaining regions of E2 are shown in grey cartoons. The boxed frames denote the regions analyzed. **b** The AS412 conformation changes upon binding with RM1-73 and CD81. AS412 residues in E2-RM1-73 and E2-CD81 are shown in orange sticks. HCs of RM Fabs and CD81 loop D are shown in magenta and lime cartoon, respectively. **c** Alignment of E2ecto structures from complexes with RM5-16, mAbs 1198, and 1382. The AS412, VR2, and VR3 of E2ecto are highlighted in orange, magenta, and dark green, respectively.

**Fig S6.**
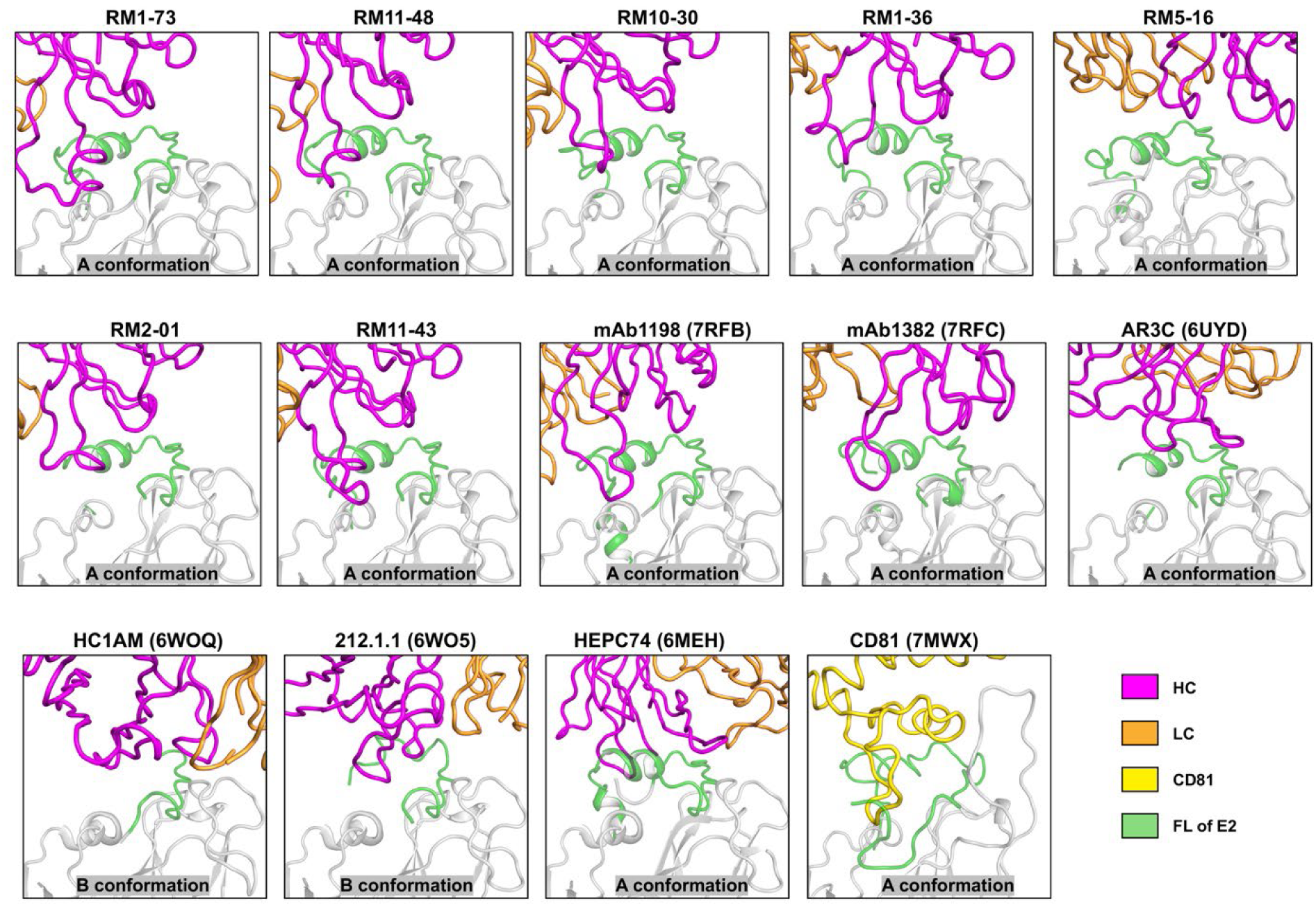
E2 FL adopts the A conformation when bound to RM AR3-targeting Abs. Close-up view of the E2 FL upon binding to AR3-targeting Abs and CD81 receptor. The HC (magenta) and LC (light orange) of the Abs are shown as ribbons. CD81 receptor is shown in a yellow ribbon. E2 proteins are shown as grey with their FL regions in green cartoon tubes.

**Fig S7.**
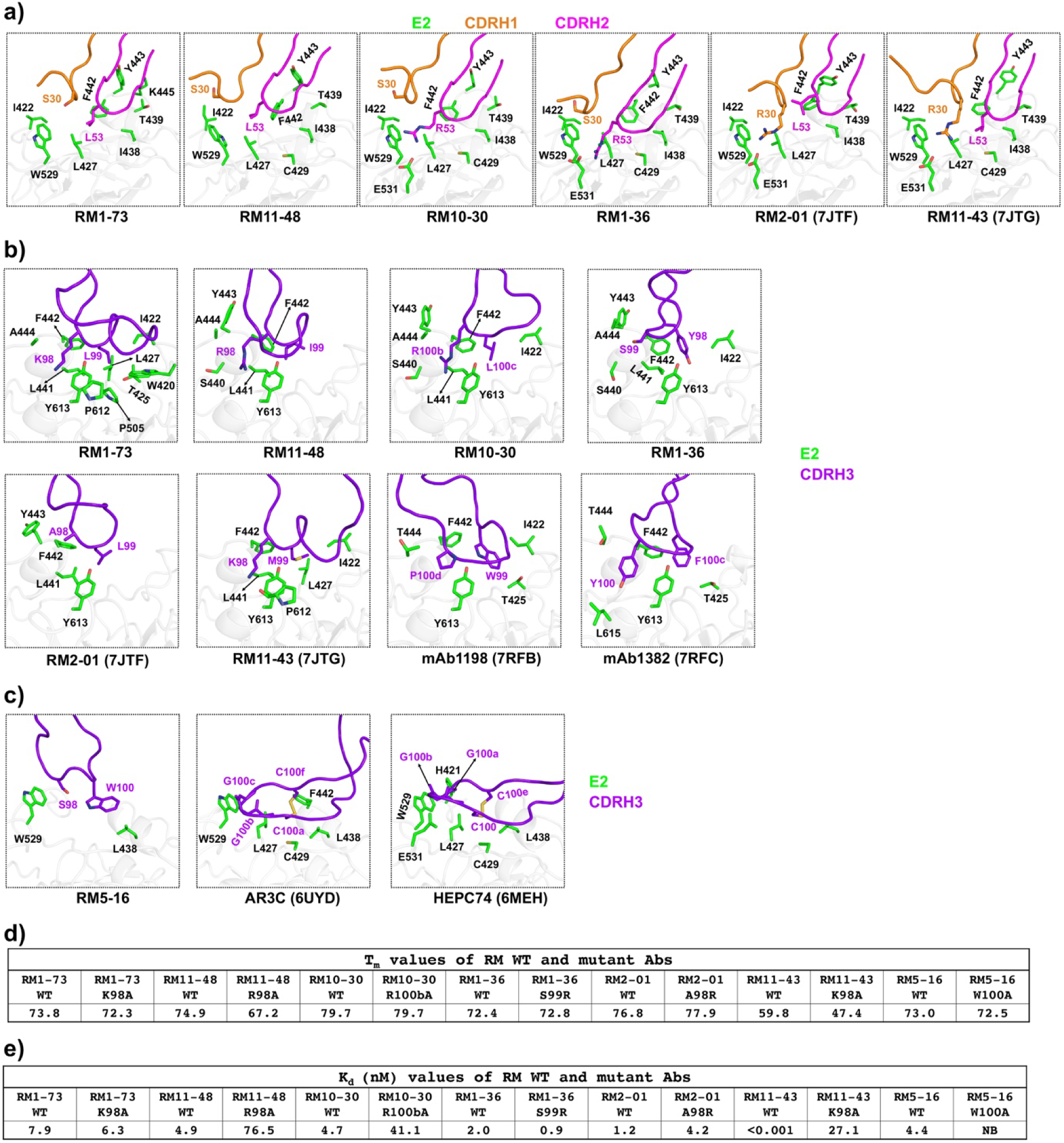
CDRH1-3 and key-residue mutational analysis of AR3-targeting Abs upon binding with E2. All complexes were superimposed based on E2 (grey cartoon). CDRs are shown as tubes, with side chains as sticks. CDRH1-3 loops are shown in orange, magenta, and purple, respectively. E2 key residues with BSA greater than 0 Å² and located within 3.9 Å of the CDRH1-3 loops are shown as green sticks. **a** Superposition of all CDRH1-2 indicating variation in CDRH1-2 residues that interact with E2. The key interacting residues of CDRH1-2 are shown in orange and magenta sticks, respectively. **b, c** Superposition of all RM CDRH3 indicates variation in CDRH3, despite a similar binding mode to E2. The key binding residues of CDRH3 are shown. **d** Comparison of melting temperature (Tm) between RM mutant and wild-type (WT) Abs. **e** Binding affinity dissociation constant (Kd) comparisons between RM mutant and WT Abs.

**Table S1.** Data collection and refinement statistics of RM unliganded Fab structures.

|  | RM5-16<br>(PDB ID: 9MS9) | RM1-73<br>(PDB ID: 9MRZ) |
| --- | --- | --- |
| **Data collection** |  |  |
| X-ray source | SSRL 12-2 | Rigaku MicroMax-007 |
| Wavelength (Å) | 0.979 | 1.542 |
| Space Group | P2<sub>1</sub>2<sub>1</sub>2<sub>1</sub> | P2<sub>1</sub> |
| Cell dimensions<br>a, b, c (Å)<br>$\alpha$, $\beta$, $\gamma$ (°) | 43.1, 70.2, 161.1<br>90, 90.0, 90 | 88.9, 73.3, 99.7<br>90, 94.2, 90 |
| Resolution (Å)<sup>a</sup> | 42.64 - 1.46 (1.51-1.46) | 44.79 - 2.03 (2.10 - 2.03) |
| R<sub>sym</sub><sup>a,b</sup><br>R<sub>pim</sub><sup>a,b</sup> | 0.15 (>1)<br>0.04 (0.48) | 0.05 (0.33)<br>0.03 (0.20) |
| Unique reflections | 85,214 | 80,829 |
| Average I/$\sigma$(I)<sup>a</sup> | 24.1 (1.6) | 26.4 (3.6) |
| CC<sub>1/2</sub><sup>a</sup> | 0.99 (0.69) | 0.99 (0.88) |
| Completeness (%)<sup>a</sup> | 99.4 (99.1) | 97.7 (93.3) |
| Redundancy<sup>a</sup> | 13.1 (13.1) | 3.8 (3.2) |
| **Refinement** |  |  |
| Resolution range (Å) | 42.64 - 1.46 (1.51 - 1.46) | 44.79 - 2.03 (2.10 - 2.03) |
| Reflections in refinement | 84,877 (8,259) | 80,796 (7,733) |
| R<sub>cryst</sub><sup>c</sup><br>R<sub>free</sub><sup>d</sup> | 0.14<br>0.17 | 0.17<br>0.19 |
| No. atoms |  |  |
| Fab | 3,285 | 6,636 |
| Ligand: SO<sub>4</sub> | 20 | 40 |
| Ligand: NHE<sup>e</sup> | 0 | 26 |
| Waters | 434 | 729 |
| Wilson B-value (Å<sup>2</sup>) | 14 | 31 |
| Average B-values (Å<sup>2</sup>) |  |  |
| Fab | 17 | 33 |
| Ligand: SO<sub>4</sub> | 41 | 71 |
| Ligand: NHE | 0 | 47 |
| Water | 31 | 41 |
| R.M.S deviations |  |  |
| Bond length (Å) | 0.008 | 0.005 |
| Bond angles (°) | 1.08 | 0.76 |
| Ramachandran (%)<sup>f</sup> |  |  |
| Favored/outlier | 97.9 (0) | 98.5 (0) |
| <sup>a</sup> Values in parentheses denote outer-shell statistics.<br><sup>b</sup> $R_{sym} = \sum_{hkl} \sum_i \vert I_{hkl,i} - \langle I_{hkl} \rangle\vert / \sum_{hkl} \sum_i I_{hkl,i}$ and $R_{pim} = \sum_{hkl} [1/(N-1)] 1/2 \sum_i \vert I_{hkl,i} - \langle I_{hkl} \rangle\vert / \sum_{hkl} \sum_i I_{hkl,i}$, where $I_{hkl,i}$ is the scaled intensity of the $i^{th}$ measurement of reflection h, k, l, $\langle I_{hkl} \rangle$ is the average intensity for that reflection, and N is the redundancy.<br><sup>c</sup> $R_{cryst} = \sum_{hkl} \vert F_o - F_c\vert / \sum_{hkl} \vert F_o\vert $, where $F_o$ and $F_c$ are the observed and calculated structure factors.<br><sup>d</sup> $R_{free}$ was calculated as for $R_{cryst}$, but on 5% of data excluded before refinement.<br><sup>e</sup> NHE is 2-(cyclohexylamino)ethanesulfonic acid (CHES).<br><sup>f</sup> Values are percentage of residues in favored and outlier regions using MolProbity <sup>71</sup>. | | |

**Table S2.**
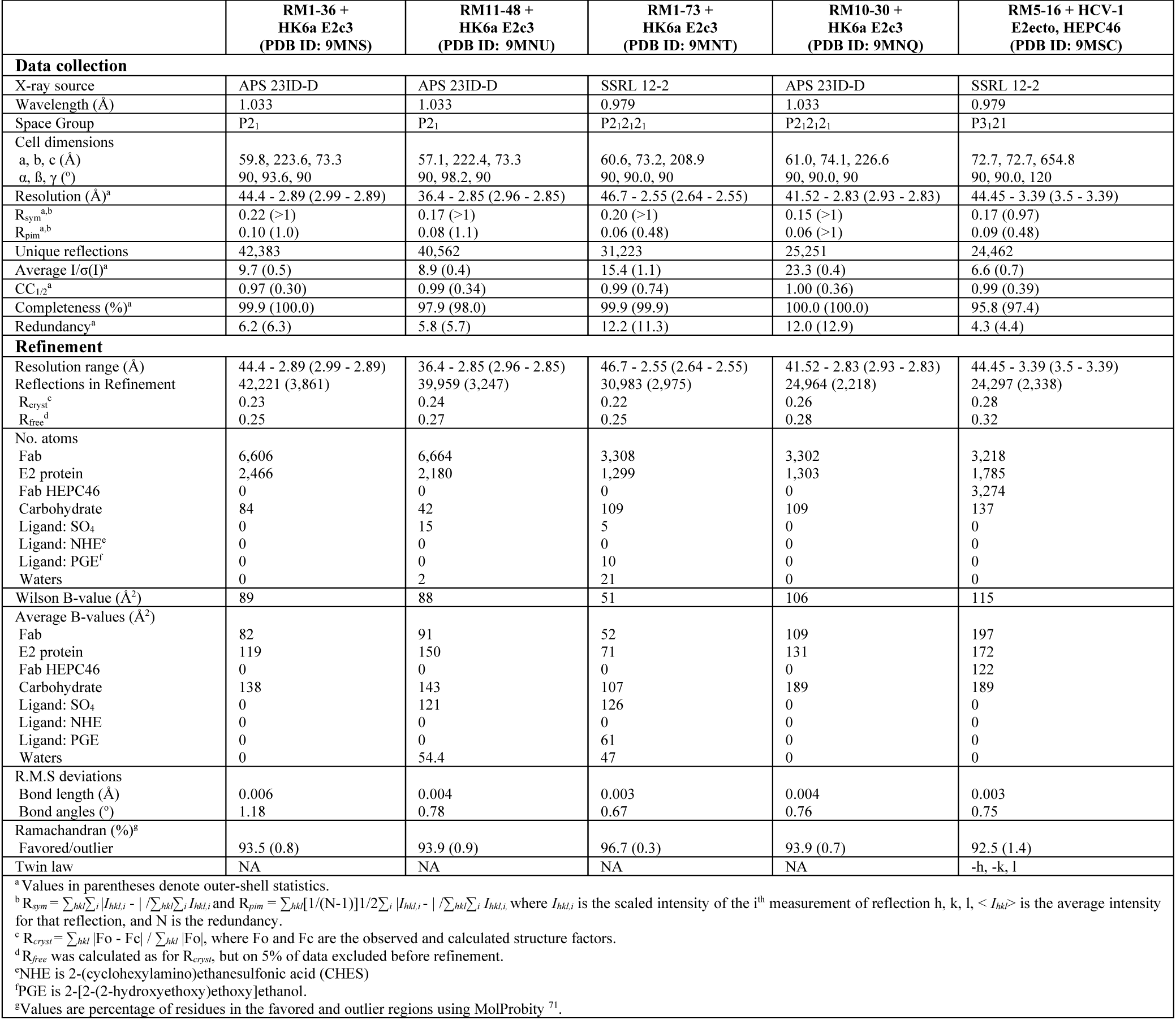
Data collection and refinement statistics of RM Fab complex structures.

**Table S3.** Hydrogen bond and charged interaction distances between Fab and E2.

| RM1-73 + E2c3 |  |  | RM11-48 + E2c3 |  |  | RM10-30 + E2c3 |  |  | RM1-36 + E2c3 |  |  | RM5-16 + E2ecto |  |  |
| --- | --- | --- | --- | --- | --- | --- | --- | --- | --- | --- | --- | --- | --- | --- |
| Fab<br>atom | E2<br>atom | Dist<br>(Å) | Fab<br>atom | E2<br>atom | Dist<br>(Å) | Fab<br>atom | E2<br>atom | Dist<br>(Å) | Fab<br>Atom | E2<br>atom | Dist<br>(Å) | Fab<br>atom | E2<br>atom | Dist<br>(Å) |
| H:E50 OE1 | Y443 OH | 2.9 | H:E50 OE1 | Y443 OH | 2.4, 2.6* | H:T28 OG1 | N423 NAG O3 | 3.0 | H:E50 OE1 | Y443 OH | 2.8, 3.0 | H:Y33 OH | E531 OE2 | 2.5 |
| H:N58 OD1 | K445 NZ | 3.4 | H:R98 NH1 | S440 O | 2.9, 2.7 | H:E50 OE1 | Y443 OH | 3.1 | H:R53 NE | L427 O | 3.9, 2.9 | H:G53 O | N532 NAG O7 | 3.0 |
| H:K98 N | F442 O | 2.8 | H:R98 NH2 | P612 O | 2.9, 3.0 | H:R53 NH2 | W529 NE1 | 3.2 | H:R53 NH1 | E531 OE2 | 2.6, 3.3 | H:T54 OG1 | E531 OE1 | 3.0 |
| H:K98 NZ | P612 O | 3.1 | H:I99 N | Y613 OH | 3.9, 3.4 | H:R53 NH1 | E531 OE2 | 3.0 | H:R53 NH2 | W529 O | 3.0, 3.2 | H:R56 NE | E531 OE2 | 3.5 |
| H:L99 N | Y613 OH | 3.2 | H:N100f ND2 | F442 O | 3.2, 3.0 | H:L100a O | F442 O | 3.0 | H:R53 NH2 | L427 O | 4.0, 3.2 | H:R56 NE | E531 OE1 | 3.6 |
| H:A100c O | W420 O | 2.8 | L:K92 NZ | N446 ND2 | 2.9, 3.7 | H:R100b NH1 | Y443 O | 3.2 | H:R53 NH2 | N428 N | 4.0, 3.3 | H:R56 NH2 | E531 OE2 | 2.8 |
| H:A100c O | W420 N | 2.9 | L:K92 O | A444 O | 3.7, 2.7 | H:R100b NH1 | L441 O | 3.2 | H:R53 NH2 | N428 OD1 | 4.8, 3.3 | H:E100a OE2 | W420 O | 3.0 |
| H:A100c O | S419 OG | 3.0 | L:N93 OD1 | N446 OD1 | 3.9, 2.4 | H:R100b NH2 | S440 O | 3.3 | H:R53 NH2 | N428 ND2 | 4.7, 3.3 | L:K53 NZ | H421 NE2 | 3.3 |
|  |  |  | L:N93 OD1 | N446 ND2 | 2.2, 3.5 | H:R100b NH1 | S440 O | 2.5 | H:Y98 N | F442 O | 3.2, 3.1 | L:N93 OD1 | N434 OD1 | 3.1 |
|  |  |  | L:N93 OD1 | A444 O | 3.1, 3.7 | H:R100d NH1 | F442 O | 2.9 | H:Y98 OH | Y613 OH | 3.1, 3.2 | L:S94 OG | N434 ND2 | 3.0 |
|  |  |  | L:N93 OD1 | N446 N | 2.7, 2.6 | L:E27 OE1 | K445 NZ | 2.7 | H:S99 N | L441 O | 3.3, 3.0 |  |  |  |
|  |  |  | L:N93 ND2 | N446 ND2 | 3.0, 3.2 |  |  |  | H:S99 OG | S440 O | 3.0, 2.8 |  |  |  |
|  |  |  |  |  |  |  |  |  | H:S99 OG | Y443 O | 3.3, 2.9 |  |  |  |
|  |  |  |  |  |  |  |  |  | H:S99 OG | L441 O | 3.5, 3.1 |  |  |  |
|  |  |  |  |  |  |  |  |  | H:S99 OG | A444 N | 3.5, 3.3 |  |  |  |
|  |  |  |  |  |  |  |  |  | H:E100 OE1 | Y613 OH | 3.0, 2.7 |  |  |  |
|  |  |  |  |  |  |  |  |  | H:Y100a OH | H421 NE2 | 2.3, NA |  |  |  |

\*: 2 distances are listed for structures with 2 complexes in the asymmetric unit NA: this residue was not modelled in one of the complexes in the asymmetric unit

**Table S4.** Van der waal contacts between Fab and E2.

| RM1-73 + E2c3 |  | RM11-48 + E2c3 |  | RM10-30 + E2c3 |  | RM1-36 + E2c3 |  | RM5-16 + E2ecto |  |
| --- | --- | --- | --- | --- | --- | --- | --- | --- | --- |
| Fab residue | E2 residue | Fab residue | E2 residue | Fab residue | E2 residue | Fab residue | E2 residue | Fab residue | E2 residue |
| H:T28 | W529 | H:T28 | N423 glycan | H:T28 | N423 glycan | H:T28 | W529 | H:Y33 | W529, E531 |
| H:I31 | I422, L427, F442, W529 | H:V31 | F442 | H:S30 | N423 glycan | H:S30 | W529 | H:Y52 | S528, W529, E531 |
| H:E50 | Y443 | H:A33 | Y443 | H:I31 | L427, F442, W529 | H:I31 | F442, W529 | H:G53 | N532 glycan |
| H:I52 | F442, Y443 | H:E50 | Y443 | H:E50 | Y443 | H:E50 | Y443 | H:T54 | N532 glycan, E531 |
| H:L53 | L427, F442 | H:I52 | F442, Y443 | H:I52 | Y443 | H:I52 | F442, Y443 | H:R56 | E531 |
| H:V54 | I438, F442 | H:L53 | L427 | H:R53 | W529 | H:R53 | L427, N428, W529, E531 | H:G97 | W529 |
| H:I56 | I438, T439, Y443 | H:V54 | I438, F442 | H:V54 | F442 | H:V54 | C429, I438, F442 | H:S98 | W529 |
| H:N58 | Y443, K445 | H:I56 | T439 | H:I56 | I438, T439, Y443 | H:L56 | T439, Y443 | H:W100 | L438 |
| H:S97 | F442 | H:R98 | S440, L441, F442, Y443, A444, Y613 | H:V58 | Y443 | H:T58 | Y443 | H: E100a | W420 |
| H:K98 | L441, F442, A444, P612, Y613 | H:I99 | L441, F442, Y613 | H:L100a | I422, L441, F442, Y613 | H:R97 | F442, Y443 | L:G30 | A439, F442 |
| H:L99 | W420, I422, T425, L427, L441, F442, P505, Y613 | H:A100b | P612 | H:R100b | S440, L441, F442, Y443, A444, Y613 | H:Y98 | I422, L441, F442, Y613 | L:Y31 | L438, F442 |
| H:V100b | W420, Y507, P612, Y613 | H:G100c | Y613 | H:R100d | F442, Y443 | H:S99 | S440, L441, F442, Y443, A444 | L:F32 | W420 |
| H:A100c | S419, W420, I422 | H:N100f | F442, Y443 | L:I2 | K445 | H:E100 | P612, Y613 | L:Q50 | W420, H421 |
| H:A100d | S419 | L:F27d | A444 | L:E27 | K445 | H:Y100a | H421, I422 | L:K53 | H421 |
| H:G100e | G418 | L:F28 | R614, L615, V622 | L:Y92 | A444, K445 | L:Y96 | Y443 | L:N93 | D431, N434 |
| L:Y32 | A444 | L:K92 | A444, N446 | L:G93 | A444, K445 |  |  |  |  |
| L:D93 | K445, N446 | L:N93 | A444, K445, N446 | L:T94 | Y443 |  |  |  |  |
| L:T94 | Y443, K445 | L:S94 | Y443, K445 |  |  |  |  |  |  |
|  |  | L:P95 | Y443, K445 |  |  |  |  |  |  |
|  |  | L:Y96 | Y443 |  |  |  |  |  |  |

**Table S5.** BLI binding of RM Abs with E2. RM Fabs are tested on various HCV isolates from different genotypes. Fabs were immobilized on human Fab-2G biosensors and E2 proteins from various isolates were tested as analytes. The fitting statistics such as K_d_, association (K_on_), K_dis_ (dissociation), and R^2^ (goodness of fit) are summarized. NB (no binding).

| HCV Genotypes | HCV Isolates | Ab ID | Kd(M) | Kon(1/Ms) | Kdis(1/s) | Full R2 |
| --- | --- | --- | --- | --- | --- | --- |
| 1a | H77 | RM11-48 | 5.65E-08 | 8.10E+03 | 4.58E-04 | 0.99 |
|  |  | RM1-73 | <1.0E-12 | 5.74E+03 | <1.0E-07 | 0.98 |
|  |  | RM10-30 | 5.48E-08 | 3.22E+03 | 1.76E-04 | 0.99 |
|  |  | RM1-36 | 3.97E-07 | 2.86E+03 | 1.13E-03 | 0.99 |
|  |  | RM5-16 | NB | NB | NB | NB |
|  | HCV-1 | RM11-48 | 4.89E-09 | 1.46E+04 | 7.16E-05 | 0.99 |
|  |  | RM1-73 | 7.88E-09 | 1.34E+04 | 1.06E-04 | 0.99 |
|  |  | RM10-30 | 4.72E-09 | 2.59E+04 | 1.22E-04 | 0.99 |
|  |  | RM1-36 | 2.01E-09 | 1.34E+04 | 2.69E-05 | 0.99 |
|  |  | RM5-16 | 4.40E-09 | 6.41E+03 | 2.82E-05 | 0.99 |
| 2a | J6 | RM11-48 | 6.06E-08 | 1.09E+05 | 6.60E-03 | 0.97 |
|  |  | RM1-73 | 3.80E-09 | 1.14E+05 | 4.31E-04 | 0.96 |
|  |  | RM10-30 | 2.58E-08 | 2.43E+05 | 6.25E-03 | 0.98 |
|  |  | RM1-36 | 2.37E-07 | 3.84E+04 | 9.09E-03 | 0.99 |
|  |  | RM5-16 | 1.37E-08 | 6.24E+04 | 8.82E-04 | 0.98 |
| 3a | S52 | RM11-48 | 2.64E-09 | 7.01E+04 | 1.85E-04 | 0.96 |
|  |  | RM1-73 | 1.82E-09 | 6.13E+04 | 1.11E-04 | 0.97 |
|  |  | RM10-30 | 5.81E-09 | 1.04E+05 | 6.06E-04 | 0.98 |
|  |  | RM1-36 | 3.30E-08 | 3.16E+04 | 1.04E-03 | 0.98 |
|  |  | RM5-16 | 1.10E-09 | 6.58E+04 | 7.21E-05 | 0.97 |
| 6a | HK6a | RM11-48 | 4.80E-09 | 1.13E+05 | 5.41E-04 | 0.99 |
|  |  | RM1-73 | 7.65E-10 | 1.33E+05 | 1.01E-04 | 0.99 |
|  |  | RM10-30 | 1.43E-09 | 2.47E+05 | 3.54E-04 | 0.99 |
|  |  | RM1-36 | 5.08E-08 | 1.02E+05 | 5.16E-03 | 0.98 |
|  |  | RM5-16 | 1.59E-08 | 1.73E+04 | 2.74E-04 | 0.99 |

**Table S6.** Crystallization conditions.

| Fab/complex | Temperature (°C) | Conditions |
| --- | --- | --- |
| RM5-16 | 20 | 0.1 M sodium acetate 4.5, 0.2 M lithium sulfate, 28 %(w/v) polyethylene glycol 8,000 |
| RM1-73 | 20 | 0.1 M CHES, pH 9.5, 0.2 M sodium chloride, 1.26 M ammonium sulfate |
| RM1-36 + HK6a E2c3 | 20 | 0.2 M tri-sodium citrate, 20 %(w/v) polyethylene glycol 3,350 |
| RM1-73 + HK6a E2c3 | 20 | 0.1 M Tris 8.5, 0.2 M lithium sulfate, 40 %(v/v) polyethylene glycol 400 |
| RM10-30 + HK6a E2c3 | 20 | 0.1 M HEPES 7.5, 0.8 M potassium dihydrogen phosphate, 0.8 M sodium dihydrogen phosphate |
| RM11-48 + HK6a E2c3 | 20 | 0.1 M CHES pH 9.5, 0.2 M sodium chloride, 1.26 M ammonium sulfate |
| RM5-16 + HCV-1 E2ecto, HEPC46 | 20 | 0.3 M magnesium chloride, 15 % (w/v) polyethylene glycol 3,350 |

